# Membrane Binding Controls the ATPase Cycle and Localization of MinD in *Bacillus subtilis*

**DOI:** 10.1101/2024.07.08.602513

**Authors:** Helge Feddersen, Charlotte Dyckmans, Marc Bramkamp

**Affiliations:** Institute for General Microbiology, Christian-Albrechts-University Kiel, Am Botanischen Garten 1-9, 24118 Kiel, Germany

**Keywords:** *B. subtilis*, Min system, cell division, ATPase, membrane binding, protein patterns, reaction diffusion, self-organization, SMLM, SMT

## Abstract

Bacteria precisely regulate the place and timing of their cell division. One of the best-understood systems for division site selection is the Min system in *Escherichia coli*. In *E. coli*, the Min system displays remarkable pole-to-pole oscillation, creating a time-averaged minimum at the cell’s geometric center, which marks the future division site. Interestingly, the Gram-positive model species *Bacillus subtilis* also encodes homologous proteins: the cell division inhibitor MinC and the Walker-ATPase MinD. However, *B. subtilis* lacks the activating protein MinE, which is essential for Min dynamics in *E. coli*. We have shown before that the *B. subtilis* Min system is highly dynamic and quickly relocalizes to active sites of division. This raised questions about how Min protein dynamics are regulated on a molecular level in *B. subtilis*. Here, we show with a combination of *in vitro* experiments and *in vivo* single-molecule imaging that the ATPase activity of *B. subtilis* MinD is activated by membrane binding. Additionally, both monomeric and dimeric MinD bind to the membrane, and binding of ATP to MinD is a prerequisite for fast membrane detachment. Single-molecule localization microscopy data confirm membrane binding of monomeric MinD variants. However, only wild type MinD enriches at cell poles and sites of ongoing division, likely due to interaction with MinJ. Monomeric MinD variants and locked dimers remain distributed along the membrane and lack the characteristic pattern formation. Single-molecule tracking data further support that MinD has a freely diffusive population, which is increased in the monomeric variants and a membrane binding defective mutant. Thus, MinD dynamics in *B. subtilis* under the tested conditions do not require any unknown protein component and can be fully explained by MinD’s binding and unbinding kinetics with the membrane. The spatial organization of MinD relies on the short-lived temporal residence of MinD dimers at the membrane.

## Introduction

Accurate selection of the division site is essential in bacteria for maintaining cell size, ensuring proper chromosome segregation, and preventing the formation of anucleate minicells. Cell division is governed by a complex protein machine known as the divisome (1–3). The spatiotemporal organization of this protein complex is directed by the dynamic arrangement of a bacterial tubulin homolog, FtsZ (4, 5). FtsZ forms treadmilling protofilaments that condense into a ring-like structure at the division site (6–8), recruiting several cell wall synthesis proteins, such as division-specific transglycosylases and transpeptidases (9, 10). Together, these proteins synthesize the bacterial peptidoglycan cell wall. To ensure that FtsZ polymerizes only at the correct location, many bacterial species employ the Min system, a conserved pathway that prevents aberrant polar septation events.

The Min system was identified through the formation of small, round, anucleate cells that divide close to the cell poles (11). The gene locus responsible for the “*min*iature” cell phenotype in *E. coli* was termed *minB* (12, 13), later shown to encode the three structural genes *minC*, *minD* and *minE* (14). MinC is the actual inhibitor of FtsZ-polymer formation (15, 16). The MinC protein contains an N- and C-terminal domain (17), both of which contribute to FtsZ inhibition. The C-terminal domain binds to the C-terminal tail of FtsZ, whereas the N-terminal domain, which also mediates MinC dimerization, binds at the interface of two FtsZ subunits (18–20). MinC-mediated inhibition of FtsZ has also been demonstrated in *B. subtilis* (21–23), suggesting a conserved mechanism across species.

The spatial localization of MinC is regulated by MinD, a Walker-type ATPase that belongs to the ParA/MinD family of proteins (21, 24, 25). These partitioning ATPases are involved in positioning cargo within bacterial cells (26, 27). While the molecular details of partitioning differ among ParA-like proteins, a common feature is their use of binding to a large intracellular surface as a spatial template (26, 27). ParA proteins typically bind non-specifically to DNA, covering the nucleoid and exploiting it as a track for cargo positioning (28). In contrast, MinD proteins bind to the membrane via an amphipathic helix (29, 30). In *E. coli*, membrane binding of MinD strictly depends on ATP binding and subsequent dimerization (24). Dimerized MinD recruits MinC and thereby activates MinC to inhibit FtsZ oligomerization. In *E. coli*, MinE further regulates MinD localization by binding to the membrane and interacting with MinD, stimulating ATP hydrolysis and triggering MinD release from the membrane. Slow nucleotide exchange in cytosolic MinD leads to an oscillatory cycle of membrane attachment and detachment, resulting in pole-to-pole oscillation of MinCD (31–33). This oscillatory behavior generates a temporal minimum of MinC at midcell, such that Z-ring assembly is restricted exclusively to this position. The oscillation can be reconstituted *in vitro* on planar membranes, demonstrating the robustness of this self-organizing system and a prime example of pattern formation. (34).

The Gram-positive model bacterium *B. subtilis* likewise encodes the *minC* and *minD* genes (35, 36), and loss of either results in the characteristic minicell phenotype. Thus, a role for the Min system analogous to that in *E. coli* has been proposed (21, 37). However, *B. subtilis* lacks a MinE homolog, and the Min proteins do not oscillate from pole to pole (21, 37). Instead, static imaging shows that MinD accumulates at the cell poles and at active division sites. Initial genetic and cell biological studies demonstrated that the polar scaffold protein DivIVA is essential for polar MinD localization (21). The interaction between DivIVA and MinD is mediated by the integral membrane protein MinJ (38, 39). MinJ is also part of the divisome and interacts with multiple division proteins (39). Loss of MinJ causes defects in divisome disassembly, as proteins such as FtsL and PBP2b remain associated with the nascent cell pole after division in Δ*minJ* cells (40). This function is supported by recent findings that MinD promotes FtsZ polymer disassembly at the cell poles, and that deletion of *minD* inhibits polar peptidoglycan remodeling (41).

We have recently shown that all components of the *B. subtilis* Min system, including DivIVA and MinJ, dynamically localize within the bacterial cell (40, 42–44). Interestingly, even the scaffold protein DivIVA, although highly conserved in many Gram-positive phyla, is not statically anchored at the poles but instead redistributes along the membrane from poles to septa (42–44). In contrast, DivIVA homologs in actinobacteria show greatly reduced mobility, consistent with their role in apical growth (43).

Despite these advances, the mechanism by which MinD activity and localization are regulated in the absence of MinE has remained unclear, raising the possibility of additional, unidentified regulatory factors or alternative modes of ATPase control. In this study, we address these long-standing questions by combining *in vitro* biochemical analyses with single-molecule imaging *in vivo*. We define the ATPase and membrane-binding properties of *B. subtilis* MinD and examine how its dynamics are shaped by interactions with MinJ and MinC. Our findings reveal an unexpected mechanism of MinD regulation that operates independently of MinE-like factors, providing a revised understanding of how the non-oscillatory *B. subtilis* Min system robustly controls cytokinesis and prevents aberrant septation events.

## Results

### Purification and gel filtration of His-MinD

Since localization and function of MinD appear to be strongly tied to its ATPase activity, the first logical step in describing and understanding details of the ATPase cycle was identification of the relevant biochemical characteristics, to eventually be able to correlate these properties to *in vivo* observations. To this end, *B. subtilis minD* was heterologously expressed in *E. coli*, yielding an N-terminally His-tagged fusion protein (Figure 1–figure supplement 1), as C-terminal MinD fusions have been shown to interfere with membrane binding (45). Since MinD is known for being membrane-associated, a well-established *E. coli* MinD purification protocol was adapted (46), where buffers were supplied with excess Mg^2+^-ADP to potentially reduce MinD self- and membrane interaction. Based on the respective western blot and gel filtration chromatogram (Figure 1–figure supplement 1 and Figure 1–figure supplement 2), His-MinD is not able to dimerise when purified in presence of excess Mg^2+^-ADP, as evidenced by a main peak at the expected sizes of the monomeric form (∼31 kDa, 16.11 ml) and the absence of a peak at the expected size of the dimeric form (∼62 kDa, 14.65 ml, Figure 1–figure supplement 2). However, Western blotting of Ni-NTA and size-exclusion chromatography (SEC) fractions indicates the presence of multiple His-tagged degradation products alongside full-length His-MinD (Figure 1–figure supplement 1). Because these fragments retain the His-tag, they co-purify during Ni-NTA and can co-elute or form non-covalent assemblies with full-length MinD, increasing the heterogeneity of the sample and likely producing the minor shoulder observed at an apparent molecular weight of ∼44 kDa (15.4 ml, Figure 1–figure supplement 2). Unfortunately, the Mg^2+^-ADP excess in the purification buffers did not seem to stop His-MinD membrane interaction, as a significant fraction of the recombinant protein were later detected to be lost in the cell pellet. However, sufficiently high His-MinD concentrations could be obtained to proceed with biochemical analysis.

### *B. subtilis* MinD displays full ATPase activity when incubated with ATP and liposomes

Being able to purify His-MinD allowed us to investigate the biochemical conditions affecting the MinD ATPase cycle next. For this purpose, we choose to use the commercially available enzyme coupled EnzChek™ phosphate assay kit (Thermo Fisher Scientific) to measure ATPase activity. The activity was determined by measuring the P_i_ production rate (in µM min^−1^) as a function of His-MinD concentration. We first tested His-MinD ATPase activity at protein concentrations between 0.5 µM and 16 µM. Under these conditions, only baseline ATPase activity below the detection limit was observed (consistent with Figure 1 D, 0 mg ml^-1^ liposomes condition). Since MinD interacts with the membrane and it was expected that the ATPase cycle is involved in controlling this interaction, we next measured ATPase activity in the presence of liposomes extruded from *E. coli* total lipids (Avanti). Remarkably, this led to a robust increase in ATPase activity of His-MinD, appearing to be linearly dependent on protein concentration (Figure 1 A & B). To be able to compare the activity level to the published *E. coli* MinD *in vitro* ATPase data (47–49), the specific activity is shown as a specific activity in nmol mg^-1^ min^-1^ (Figure 1 B). Interestingly, activity levels of recombinant *B. subtilis* MinD were found to be on par with fully active *E. coli* MinD in conjunction with MinE and phospholipids (47). Moreover, the ATPase activity did not show cooperative behavior compared to *E. coli* MinD at lower concentrations (47). We further measured that ATP hydrolysis can be titrated with both ATP and liposomes and is described by Michaelis-Menten kinetics (Figure 1 C & D). Again, the hereby-obtained V_max_ values (19.50 nmol mg^-1^ min^-1^ and 22.05 nmol mg^-1^ min^-1^, respectively) indicate similar ATPase activity levels between MinD from *E. coli* and *B. subtilis*.

**Figure 1:**
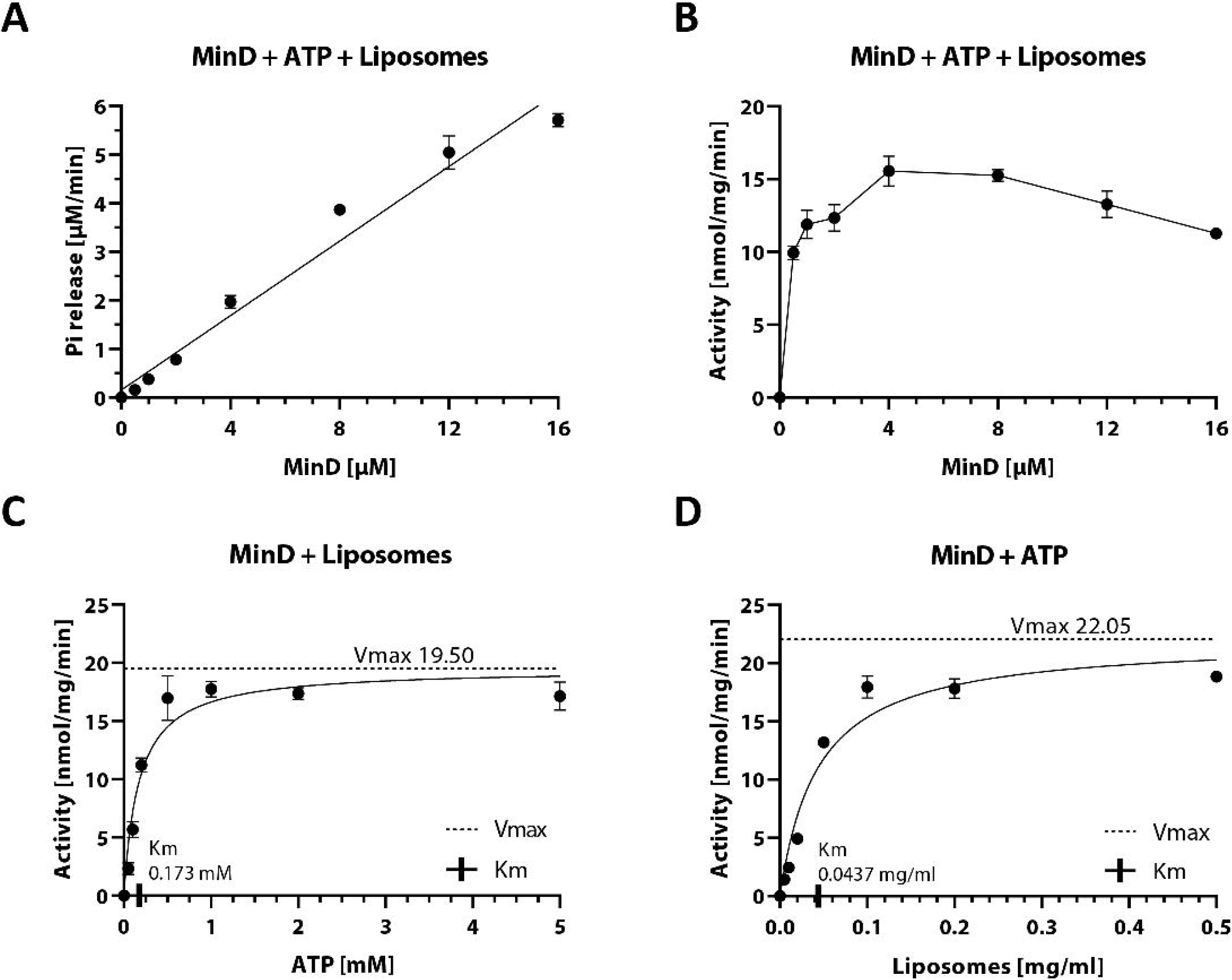
Biochemical analysis of *B. subtilis* MinD ATPase cycle. (A) Phosphate release plotted against different MinD concentrations, fitted with a simple linear regression (R² = 0.97). Phosphate release was measured using the EnzChek™ phosphate assay kit. Samples contained 0.2 mg ml^-1^ liposomes and were pre-incubated for 10 minutes before addition of 2 mM Mg^2+^-ATP; n ≥ 3. (B) Specific activity of the MinD ATPase from (A) measured as a function of MinD concentration. (C) Specific activity of MinD measured as a function of ATP concentration, determined as in (B) with fixed MinD concentration of 10 µM. Fitting the Michaelis–Menten equation (black lines) gives *k*_cat_ = 36.27 h^−1^, V_max_ = 19.50 nmol mg^-1^ min^-1^ and *K_M_* = 0.173 mM. (D) Specific activity of MinD measured as a function of liposome concentration, determined as in (B) with fixed MinD concentration of 6 µM. Fitting the Michaelis–Menten equation (black lines) gives *k*_cat_ = 41.01 h^−1^, V_max_ = 22.05 nmol mg^-1^ min^-1^ and *K_M_* = 0.0437 mg ml^-1^.

### MinC and the PDZ domain of MinJ have different effects on MinD ATPase activity

Even though ATPase activity levels were similar, proteins that directly interact with MinD could potentially modulate this activity *in vivo* as seen for MinD:MinE in *E. coli* (47–49). To get the complete picture of MinD activity, investigating the potential effects of both known *B. subtilis* MinD interactors MinC and MinJ was therefore necessary. For this purpose, both proteins were subjected to purification attempts. While purification of the cytosolic His-MinC could be successfully established using affinity chromatography followed by SEC (Figure 2–figure supplement 1), the intermembrane protein His-MinJ proved more challenging. Since several different purification attempts of the full-length protein yielded no stable product, a truncated form containing only the soluble C-terminal PDZ domain of MinJ fused to an N-terminal His-tag (His-PDZ) via a flexible linker was expressed instead (Figure 2–figure supplement 2). PDZ domains are often found in membrane-associated proteins and known to selectively interact with the C-terminus of partner peptides or proteins to sequester them to highly specific sites of the cell, both in pro- and eukaryotes (50). Although His-PDZ could be isolated via Ni-NTA affinity chromatography, it was unstable during SEC and consistently lost from the column, precluding further purification by this method. Hence, further experiments were performed using His-PDZ directly after affinity chromatography.

To test the effect of His-MinC and His-PDZ on His-MinD ATPase activity, the previously established enzymatic assays were repeated in the presence of either His-MinC or His-PDZ (Figure 2). Addition of His-MinC resulted in a very small but statistically significant decrease in MinD ATPase activity (∼5%). Given the small sample size this effect should be interpreted cautiously. However, it could be caused by the experimental set-up, since the high amounts of MinC (6 µM) that have been used might sequester a small fraction of MinD away from membrane surfaces that are required for efficient ATP hydrolysis, due to direct interaction of MinC and MinD.

In contrast, addition of His-PDZ led to a slight but reproducible increase in MinD ATPase activity, which prompted us to repeat the experiment at different His-PDZ concentrations (Figure 2). All His-PDZ concentrations significantly increased His-MinD ATPase activity compared to the control (1 µM His-PDZ = 25% ± 6.8%, 3 µM His-PDZ = 37 ± 12.5 %; 5 µM His-PDZ = 22% ± 2.8 %). We speculate that the PDZ domain may promote MinD:MinD encounters, facilitating dimerization rather than directly enhancing catalytic turnover. Because only the soluble PDZ domain could be purified, it remains unclear how full-length, membrane-anchored MinJ would influence MinD activity in vivo. The magnitude of the effect might differ when the PDZ domain is present in its native membrane-associated context.

**Figure 2:**
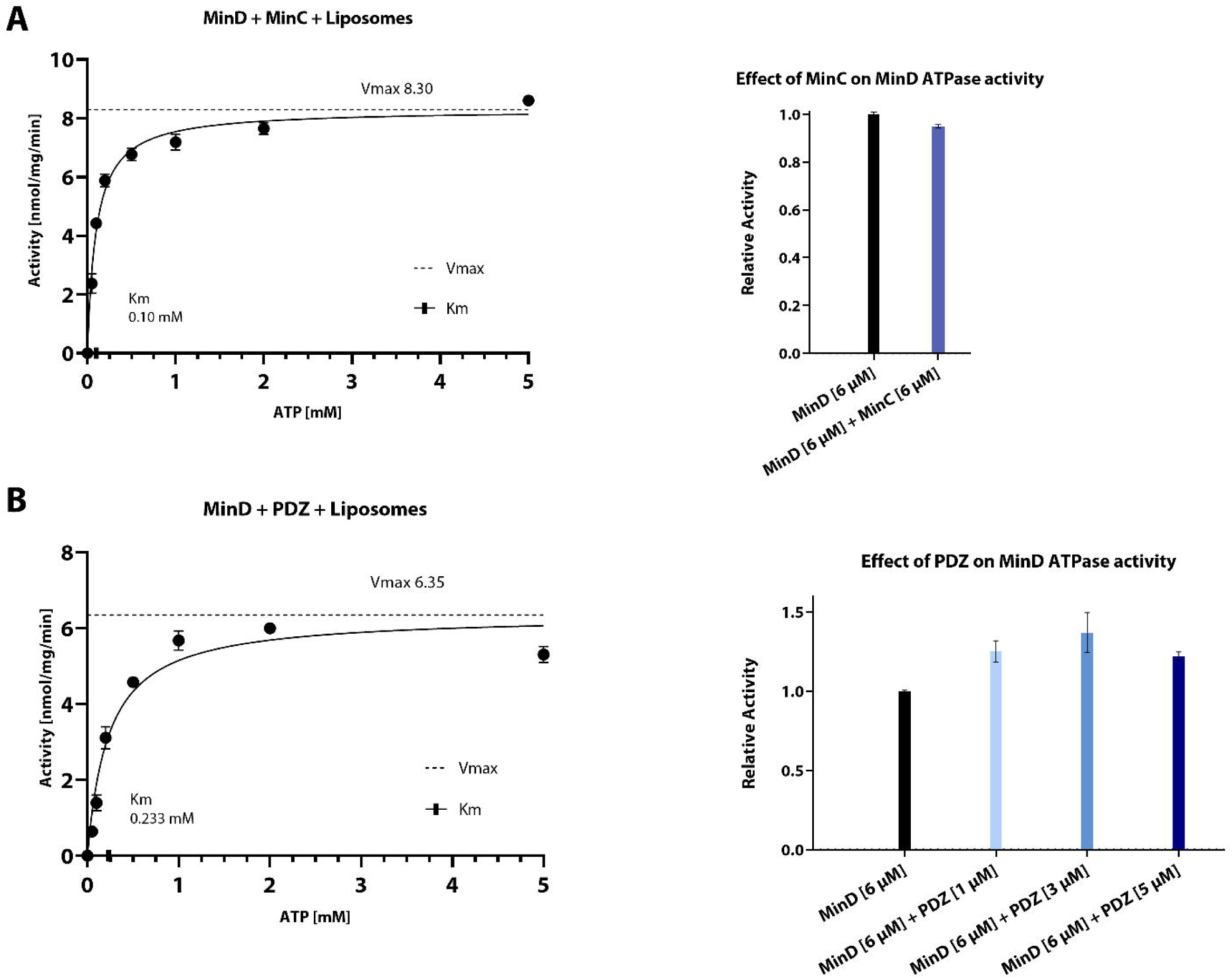
Biochemical analysis of effect of His-MinC or His-PDZ on MinD ATPase activity. (A) Left: Specific activity of MinD measured as a function of ATP concentration. Phosphate release was measured using the EnzChek™ phosphate assay kit. Samples contained 6 µM His-MinD, 6 µM His-MinC and 0.2 mg ml^-1^ liposomes and were pre-incubated for 10 minutes before addition of 2mM Mg^2+^-ATP; n ≥ 3. Fitting the Michaelis–Menten equation (black lines) gives *k*_cat_ = 15.77 h^−1^, V_max_ = 8.30 nmol mg^-1^ min^-1^ and *K_M_* = 0.10 mM. Right: Relative activity of 6 µM His-MinD in the presence of 6 µM His-MinC (blue) compared to an internal control of 6 µM His-MinD (black, relative activity = 1). Whiskers represent SD. Specific activity of only His-MinD was 8.61 [nmol mg^-1^ min^-1^] (SD = 0.03, n=2). Relative activity of 6 µM His-MinD in the presence of 6 µM His-MinC was 0.95 with SD = 0.071 (n = 3). Significance was assessed by performing Welch’s t-test on the raw data. p < 0.05. (B) Left: Specific activity of MinD measured as a function of ATP concentration. Phosphate release was measured using the EnzChek™ phosphate assay kit. Samples contained 6 µM His-MinD, 5 µM His-PDZ and 0.2 mg ml^-1^ liposomes and were pre-incubated for 10 minutes before addition of 2mM Mg^2+^-ATP n ≥ 3. Fitting the Michaelis–Menten equation (black lines) gives *k*_cat_ = 12.07 h^−1^, V_max_ = 6.35 nmol mg^-1^ min^-1^ and *K_M_* = 0.233 mM. Right: Relative activity of 6 μM His-MinD in the presence of different concentrations of His-PDZ (blue) compared to only 6 μM His-MinD (black, relative activity = 1). Whiskers represent SD. Specific activity of an internal control of His-MinD was 4.60 nmol mg^-1^ min^-1^ (SD = 0.03, n=2). Relative activities of His-MinD in the presence of 1 µM, 3 µM and 5 µM His-PDZ were 1.22, 1.37 and 1.25 with SD = 0.07, 0.13 and 0.03, respectively (n = 3 each). Significance was assessed by performing Welch’s t-test on raw data of all samples versus control. p < 0.05 for all PDZ concentrations.

### MinD ATPase activity is triggered by membrane binding

To determine whether interaction with the membrane is sufficient to stimulate MinD ATPase activity, a MinD mutant that is defective in membrane binding was required. MinD interacts with the membrane through an amphipathic α-helix located at the N-terminus, thus being the membrane targeting sequence (MTS) (Figure 3 A). Exchanging the hydrophobic isoleucine residue in the apolar core of the MTS with a charged glutamate (I260E) has been shown to abolish membrane binding of MinD by perturbing the amphipathic properties, demonstrated through fluorescence microscopy (51). Accordingly, a His-MinD-I260E variant was constructed and purified via the established purification protocol. The yield of the purification was extensively higher compared to the wild type protein (Figure 4–figure supplement 1), hinting towards a successful abolishment of membrane interaction.

**Figure 3:**
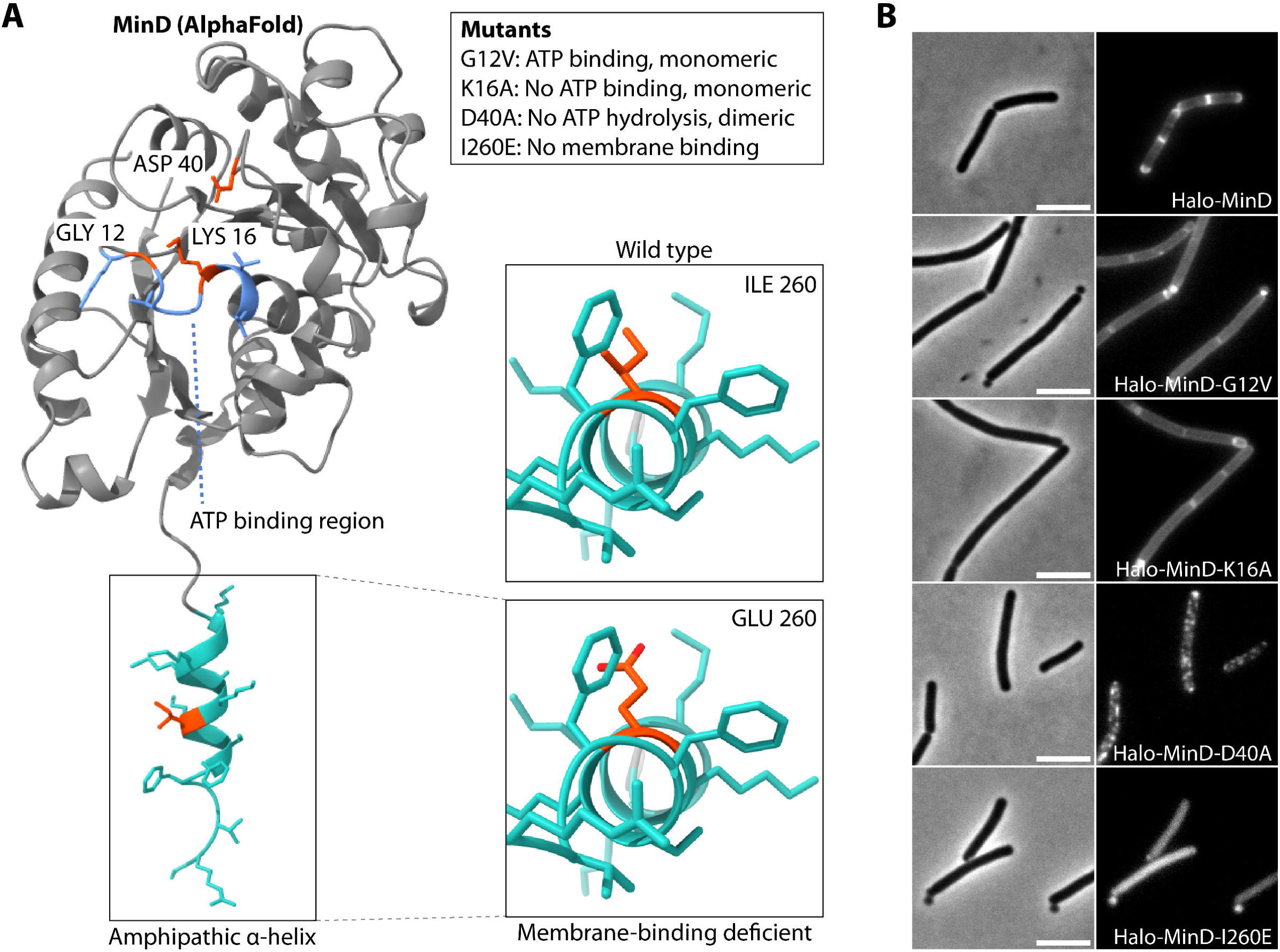
Cartoon of MinD AlphaFold model with highlighted key amino acids and the effects of their mutagenesis. (A) Left: Rendering of MinD AlphaFold model with ATP binding region highlighted in light blue, membrane targeting sequence (MTS) in turquoise and key residues in orange; Box: explanation of mutagenesis effects. Right: Close-up on hydrophobic region of amphipathic helix.(B) Representative wide-field microscopy images of *B. subtilis* expressing indicated Halo-MinD variants as the only MinD copy from the native locus, stained with TMR ligand. Left: Phase contrast; scale bar 5 µm. Right: fluorescence.

We next repeated ATPase activity assays with the purified, recombinant I260E mutant, and remarkably, activity remained below the detection limit of the assay, while a positive control of His-MinD wild type protein displayed regular activity (Figure 4). In summary, these data collectively suggest that membrane binding is necessary and the key trigger for ATP hydrolysis of *B. subtilis* MinD, without the apparent need for another stimulus.

**Figure 4:**
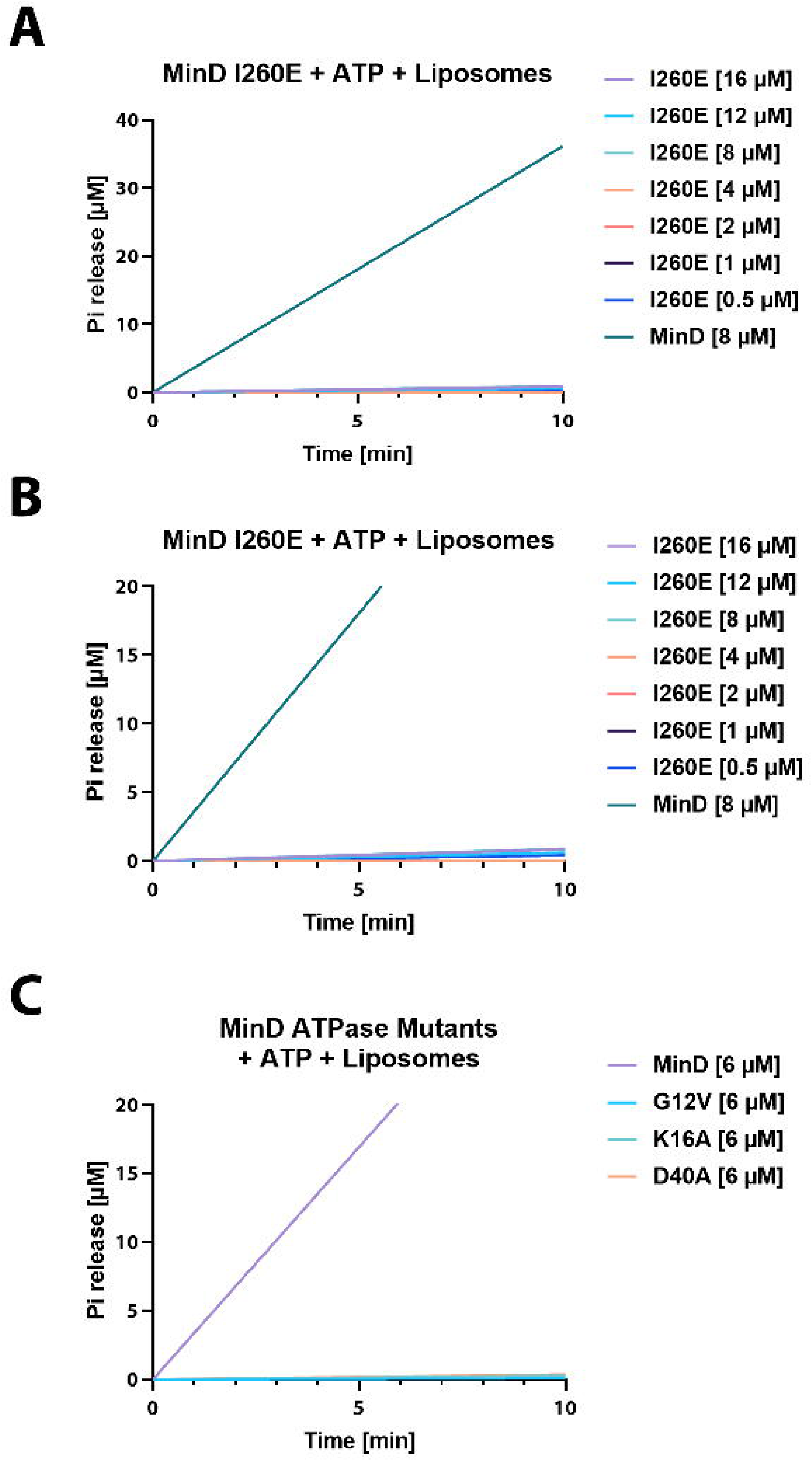
His-MinD mutants are catalytically inactive in ATP hydrolysis assay. (A) Phosphate release plotted against different His-MinD-I260E concentrations and His-MinD as control, fitted with a simple linear regression. Phosphate release was measured using the EnzChek™ phosphate assay kit. Samples contain 0.2 mg ml^-1^ liposomes and were pre-incubated for 10 minutes before addition of 2 mM Mg^2+^-ATP. (B) Same as (A) but with different y-axis scaling for better visualization of His-MinD-I260E baseline activity. (C) Same as (B) but comparing His-MinD mutants G12V, K16A and D40A to His-MinD at 6 µM concentrations, respectively.

Next, we wanted to compare *in vivo* and *in vitro* data. Therefore, a *B. subtilis* strain for microscopy was constructed, where the native copy of MinD was replaced by a HaloTag-MinD fusion protein (Figure 3 B, HaloTag will henceforth be referred to as Halo). We then introduced the I260E mutation, resulting in a strain producing Halo-MinD-I260E as the only copy. Microscopic analysis of a Halo-MinD-I260E with TMR dye confirmed drastically reduced membrane binding through its almost exclusive cytosolic localization (Figure 3 B). Furthermore, cell length and the number of minicells increased drastically, comparable to a *minD* knockout (Table 1).

**Table 1:** Measurements of cell length and minicell formation of *B. subtilis* strains.

| Strain | Relevant Genotype | Length ± SD ($\mu\text{m}$) | Minicells (%) | no. cells counted |
| --- | --- | --- | --- | --- |
| 168 | Wild type | $3.32 \pm 0.91$ | 0.20 | 610 |
| BHF077 | $\Delta\text{minD}$ | $5.80 \pm 2.10$ | 33.53 | 865 |
| BHF078 | Halo-MinD | $4.06 \pm 0.99$ | 7.44 | 430 |
| BHF079 | Halo-MinD (G12V) | $6.13 \pm 2.13$ | 34.54 | 443 |
| BHF080 | Halo-MinD (K16A) | $5.44 \pm 1.84$ | 38.45 | 489 |
| BHF081 | Halo-MinD (D40A) | $5.67 \pm 2.09$ | 33.70 | 454 |
| BHF082 | Halo-MinD (I260E) | $5.38 \pm 1.68$ | 33.01 | 721 |

### Construction of catalytic MinD mutants

To determine the role of ATP-mediated dimerization in membrane binding, we explored further mutations that affect the ATP binding region and thus the dimerization process. Due to the conserved nature of Walker-type MinD/ParA-like ATPases, residues involved in catalytic activity have been well characterized (52–54). We therefore constructed expression plasmids encoding three different MinD variants (illustrated in Figure 3 A): This first mutation, MinD-G12V, produces a steric clash within the active site, so that ATP can still be bound, but dimers cannot form (52, 53). The second mutation, K16A, completely disrupts ATP binding and hence results in monomeric protein (52). The third variant encodes the mutation D40A, which should interfere with ATP hydrolysis (53, 54). Therefore, it was expected that MinD-D40A forms a trapped nucleotide sandwich dimer similar to Soj or *E. coli* MinD (53, 54). All MinD variants were successfully expressed and purified and resulted in similar protein yields when compared to the wild type protein (Figure 1–figure supplement 1, Figure 4–figure supplement 2, Figure 4–figure supplement 3 and Figure 4–figure supplement 4). When evaluating the gel filtration chromatograms, His-MinD-G12V elute in a single peak, which can be interpreted as a monomer when compared to the wild type His-MinD elution peak (Figure 1–figure supplement 2). His-MinD-D40A on the other hand also eluted in a single peak, albeit being wider and eluting earlier, indicating a mainly dimeric conformation. It eluted between the expected dimer and monomer positions during SEC as a wide peak (Figure 1–figure supplement 2). As His-MinD-D40A cannot hydrolyze ATP, it should be dimeric and likely also form small oligomers or transient multimers. A mixture of monomer/dimer/oligomer likely produced the broad, left-shifted peak whose center is between monomer and dimer. Only His-MinD-K16A reproducibly formed a rather unexpected shoulder in its main elution peak, which could hint towards further interaction with the gel filtration matrix or other potential self-interactions. All of the recombinant His-MinD mutants displayed negligible ATPase activity when tested using the EnzChek phosphate assay kit (Figure 4 C). This was expected, as all of these MinD variants (G12V, K16A and D40A) were mutated in residues which significantly alter the ATP binding site. In all cases, activity levels remained consistently near the detection limit, precluding reliable quantitative interpretation.

To investigate *in vivo* effects of these variants, the same mutations were introduced into the strain expressing Halo-MinD in *B. subtilis*, respectively, and subsequently imaged via wide-field microscopy (Figure 3 B). All strains producing MinD mutants displayed an increased cell length and minicell production comparable to Δ*minD* (Table 1), as it was expected in cells containing dysfunctional Min proteins. Surprisingly, both monomeric MinD variants (G12V and K16A) were still observed to be membrane associated, however resulting in the apparent loss of the polar-septal MinD localization (Figure 3 B). Finally, the locked dimer D40A mutant appeared to be mainly membrane associated, thereby forming large foci indicating accumulation of MinD, but much less dispersed when compared to G12V or K16A (Figure 3 B). In summary, the time-resolved MinD gradient appears to be lost in all MinD mutants, including D40A. This suggests that the ATPase cycle and thus quick cycling of MinD is indeed necessary for formation of the observed dynamic gradient, as it has been previously suggested (44).

### Monomeric MinD binds the membrane with high efficiency

Membrane binding appeared to be a key factor of the ATPase cycle and since the microscopy-data indicated binding of MinD monomers to the membrane, we next aimed to determine biochemically, if the respective MinD variants are capable of binding to membrane and if so, to characterize their respective binding kinetics. To this end, we employed Bio-layer interferometry (BLI), a technique capable of quantifying binding strength and protein interaction, as well as identifying reaction kinetics. Liposomes were immobilized on a biosensor to create a membrane-like surface for subsequent MinD binding assays. After successful saturation of the sensor with liposomes, MinD-liposome association and dissociation was measured, a scheme of the experimental set-up is depicted in Figure 5 A. As a negative control for unspecific binding, MinD-sensor interaction was tested on empty SA sensors. Furthermore, unspecific binding to liposome-saturated sensors was tested using bovine serum albumin (BSA). Both resulted in negligible, diffuse signal (data not shown).

Kinetics of His-MinD binding was tested first, using different protein concentrations between 1 µM and 16 µM in presence of ATP. Binding to the sensor-bound liposomes scaled with increasing protein concentration, resulting in an equilibrium dissociation constant *K*_D_ = 7.17 µM and an association rate constant *k*_on_ = 3.03 x 10^3^ M^-1^ s^-1^ as a reference point for the wild type protein (Figure 5 B, Table 2). Unsurprisingly, His-MinD-I260E showed a more than 10-fold reduced affinity to the membrane with a *K*_D_ = 74.36 µM (Table 2), which is also apparent in the binding graph (Figure 5 B). The strong increase in *K*_D_ is based on the over 16-fold lowered association *k*_on_ = 0.18 x 10^3^ M^-1^ s^-1^, while the dissociation *k*_off_ = 13.45 x 10^-3^ s^-1^ was also slightly lower when compared to wild type MinD. Next, both monomeric mutants G12V and K16A were probed, and to our surprise, capable of membrane interaction (Figure 5 C & D). This is in stark contrast to *E. coli* MinD, where dimerization needs to precede membrane binding to achieve sufficient affinity (29, 30, 48, 54, 55). Similar to I260E, the G12V mutant displayed an around 10-fold reduced membrane affinity equilibrium with *K*_D_ = 75.38 µM (Table 2). However, G12V only exhibited a ∼5 fold reduced binding affinity with *k*_on_ = 0.63 x 10^3^ M^-1^ s^-1^ when compared to the wild type MinD, while the *k*_off_ = 47.8 x 10^-3^ s^-1^ represents the fastest dissociation rate constant between all MinD variants and more than 2-fold increase compared to wild type MinD (Table 2). This indicates a higher turnover and thus a quicker release of the protein from the membrane. Markedly different from this, K16A displayed a *K*_D_ of 1.36 µM, but not much lower binding rates (*k*_on_ = 2.03 x 10^3^ M^-1^ s^-1^, Table 2 & Figure 5 C). Instead, the low *K*_D_ can be mostly attributed to the reduced dissociation (*k*_off_ = 2.77 x 10^-3^ s^-1^), implying a much longer retention of His-MinD-K16A at the membrane (Figure 5 D). Since G12V can bind ATP while K16A cannot, these findings reveal that presence of ATP in the binding pocket affects association and dissociation of MinD.

**Figure 5:**
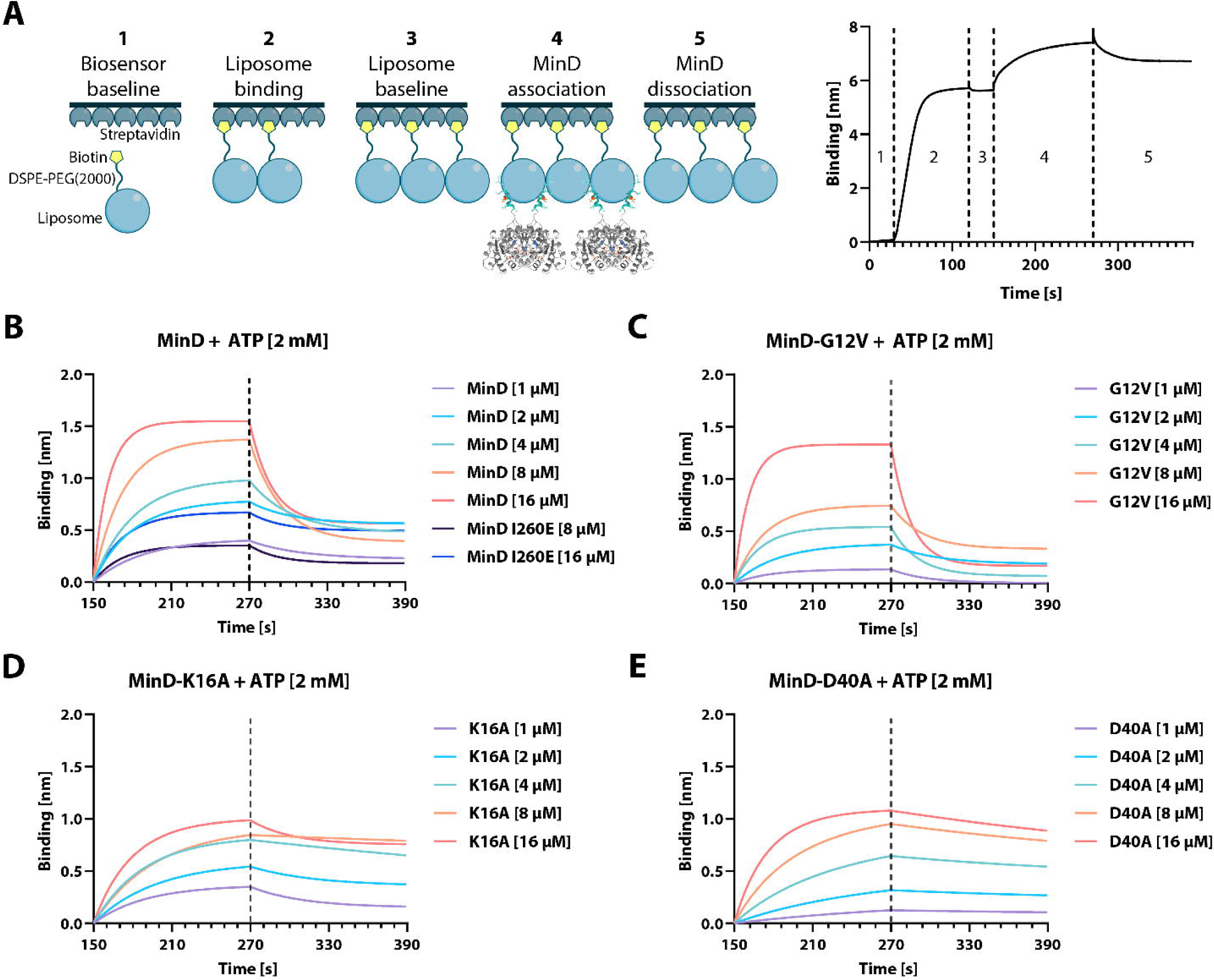
BLI analysis of His-MinD and different mutants. (A) Left: Cartoon representation of the different BLI steps, starting with (1) establishing a baseline in protein buffer, (2) binding of liposomes through the biotinylated DSPE-PEG(2000) phospholipid, (3) establishing a new baseline, (4) binding of MinD and finally (5) dissociation of MinD. Right: Exemplary graph resulting from sensor readouts of steps (1) - (5). (B) MinD binding plotted against time through association and dissociation phase (steps 4-5) at different protein concentrations, including the His-MinD-I260E mutant. (C) – (E) Same as (B) using the indicated respective mutant of His-MinD (G12V, K16A and D40A).

**Table 2:** Kinetic constants of His-MinD and variants obtained via BLI.

| Protein | $K_D$ [$\mu\text{M}$] | $k_{on}$ [$\text{M}^{-1} \text{s}^{-1}$] | $k_{on \text{ error}}$ | $k_{off}$ [$\text{s}^{-1}$] | $k_{off \text{ error}}$ | $\chi^2$ | $R^2$ |
| --- | --- | --- | --- | --- | --- | --- | --- |
| MinD | 7.17 | $3.03 \times 10^3$ | $0.13 \times 10^3$ | $21.7 \times 10^{-3}$ | $0.33 \times 10^{-3}$ | 16.07 | 0.95 |
| MinD-G12V | 75.38 | $0.63 \times 10^3$ | $0.19 \times 10^3$ | $47.8 \times 10^{-3}$ | $1.13 \times 10^{-3}$ | 13.89 | 0.95 |
| MinD-K16A | 1.36 | $2.03 \times 10^3$ | $0.062 \times 10^3$ | $2.77 \times 10^{-3}$ | $0.088 \times 10^{-3}$ | 6.97 | 0.95 |
| MinD-D40A | 0.74 | $2.39 \times 10^3$ | $0.053 \times 10^3$ | $1.76 \times 10^{-3}$ | $0.069 \times 10^{-3}$ | 6.01 | 0.98 |
| MinD-I260E | 74.36 | $0.18 \times 10^3$ | $0.19 \times 10^3$ | $13.45 \times 10^{-3}$ | $0.38 \times 10^{-3}$ | 3.61 | 0.95 |

$K_D$ = equilibrium dissociation constant; $k_{on}$ = association rate constant; $k_{off}$ = dissociation
rate constant

Last, the His-MinD-D40A mutant protein was probed, resulting in an almost 10-fold reduced dissociation equilibrium *K*_D_ = 0.74 µM (Table 2, Figure 5 E). Here, the strongly reduced membrane dissociation (*k*_off_ = 1.76 x 10^-3^ s^-1^) causes the drastic change in *K*_D_, which we expected for the trapped dimer, as it could position the two amphipathic helices into a more optimal orientation of the apolar residues towards the membrane, resulting in stronger binding and no capability of hydrolyzing ATP and thus relieving the dimer (51, 53, 54).

In summary, these BLI data demonstrate that *B. subtilis* MinD is capable of binding membrane in both monomeric and dimeric form, while the presence of ATP in the binding pocket appears to affect both membrane binding and dissociation.

### MinC and the PDZ-domain of MinJ have no significant impact on MinD membrane kinetics

*In vivo*, the integral membrane protein MinJ recruits MinD to poles and septum, which in turn recruits MinC to form a functional complex that inhibits FtsZ-ring assembly. However, the mechanistic details of how MinJ influences membrane association of MinD remain unclear. Does MinJ actively recruit MinD from the cytoplasm, or does it primarily sequester MinD already bound to the membrane? Furthermore, does MinJ promote MinD dimerization or oligomerization, potentially enhancing ATPase activity and triggering membrane dissociation? Similarly, MinC binding to MinD may influence the stability or turnover of MinD at the membrane, yet its impact on association/dissociation kinetics is unknown. To address these questions directly, we employed BLI to test the effect of His-MinC and His-PDZ on His-MinD membrane kinetics.

After validation of the experimental setup (see Material and Methods), we next tested how His-MinC and His-PDZ affect MinD-liposome binding kinetics by subjecting a liposome coated sensors to solutions containing 8 µM His-MinD and varying concentrations [1, 2, 4, 8 and 16 µM] of His-MinC or His-PDZ, respectively (Figure 5–figure supplement 1, Table 3). MinD controls appear twice in the table, because the experiments were performed using two independent purification batches. The corresponding His-MinC and His-PDZ samples were each paired with the His-MinD batch they contained. Due to batch-specific differences in His-MinD stability and degradation, we did not directly compare measurements across batches. Interestingly, neither His-MinC nor the His-PDZ domain significantly altered the membrane binding kinetics of His-MinD in BLI experiments. The extracted *K*_D_, *k_on_*, and *k_off_* values differed only marginally between conditions (Table 3), and these apparent shifts fell within the fitting uncertainty and typical inter-run variability of BLI. This contrasts with the slight increase in His-MinD ATPase activity observed in the presence of His-PDZ (Figure 2), which had led to the hypothesis that PDZ might promote MinD oligomerization at the membrane, enhancing ATP hydrolysis. However, the discrepancy between the two systems and the high lipid concentration (2 mg/ml) in the ATPase assay versus a liposome-coated sensor in BLI may account for the different outcomes. In the BLI experiment, limited surface area and distinct spatial organization of liposomes may restrict the ability of the soluble PDZ domain to effectively cluster MinD, unlike full-length MinJ, which is membrane-anchored via its transmembrane helices. Thus, while the PDZ domain alone appears insufficient to modulate membrane binding kinetics of MinD, it remains plausible that the full-length MinJ could exert a more pronounced effect through spatial coordination at the membrane.

**Table 3:** Kinetic constants of His-MinD in combination with His-MinC or His-PDZ.

| Protein | $K_D$ [$\mu\text{M}$] | $k_{on}$ [$\text{M}^{-1} \text{s}^{-1}$] | $k_{on \text{ error}}$ | $k_{off}$ [$\text{s}^{-1}$] | $k_{off \text{ error}}$ | $\chi^2$ | $R^2$ |
| --- | --- | --- | --- | --- | --- | --- | --- |
| **MinD\*** | 31.7 | $0.27 \times 10^3$ | $0.21 \times 10^3$ | $8.65 \times 10^{-3}$ | $0.26 \times 10^{-3}$ | 4.10 | 0.957 |
| **+ MinC** | 33.4 | $0.35 \times 10^3$ | $0.084 \times 10^3$ | $11.7 \times 10^{-3}$ | $0.16 \times 10^{-3}$ | 8.07 | 0.979 |
| **MinD\*** | 51.1 | $0.34 \times 10^3$ | $0.12 \times 10^3$ | $17.5 \times 10^{-3}$ | $0.24 \times 10^{-3}$ | 2.40 | 0.986 |
| **+ PDZ** | 47.5 | $0.35 \times 10^3$ | $0.14 \times 10^3$ | $16.6 \times 10^{-3}$ | $0.25 \times 10^{-3}$ | 8.98 | 0.973 |

### Single-molecule localization microscopy confirms *in vitro* data

To validate the biochemical *in vitro* results, we decided to utilize single-molecule localization microscopy (SMLM) with focus on single-molecule tracking (SMT), a technique allowing to observe and evaluate MinD dynamics *in vivo* inside *B. subtilis* cells. We began SMLM analysis by plotting a heat-map of the intracellular localization of all individually recorded Halo-MinD molecules as well as the mutated variants, respectively, on normalized cells (Figure 6 A). This representation did not only provide a robust internal control for the imaging and post-processing process for the well characterized localization of MinD, but also revealed systemic differences in localization caused by the respective mutations. The first obvious difference is reduced MinD enrichment at the cell center and poles in the G12V or K16A variants. (Figure 6 A). Here, the membrane-bound MinD fraction seems to be much more evenly distributed along the membrane, similar to what we observed during wide-field microscopy (Figure 3 B). This result does not only confirm the ability of MinD monomers to bind the membrane (Figure 5 C & D), but also fuels the speculation that MinJ might be unable to recruit MinD in its monomeric form, so that polar and septal recruitment by MinJ requires dimeric MinD. In combination with the information obtained from BLI results (Figure 5), this could mean that the membrane itself serves as proxy for MinD dimerisation and subsequent recruitment, quickly followed by ATP hydrolysis and membrane detachment of MinD. In contrast, but in agreement with wide-field imaging (Figure 3 B), the Halo-MinD-D40A mutant does appear to frequently form foci and larger local accumulations, with little protein in the cytosolic fraction (Figure 6 A). This, again, could be caused by local MinJ interaction and recruitment in combination with the missing capability of membrane detachment or with increased lateral MinD interactions (56). Finally, the I260E mutant appears evenly distributed throughout the cell (Figure 6 A), as it was expected.

Independently, all protein trajectories were subjected to a stationary localization analysis (SLA) (Figure 6 B). This was done by first grouping all trajectories into an either mobile or static population by testing if the molecule left a circle with a diameter of 97 nm (equaling the pixel size of the imaging setup) within at least five consecutive frames. Trajectories that were classified as mobile were further sub-grouped into mixed and free trajectories, where only free trajectories never presented a confinement event (57). In agreement with previous data, the D40A mutant displayed the highest fraction of static tracks (Figure 6 B), and by far the lowest proportion of truly free trajectories (48.5 %). In contrast, more than 90% of Halo-MinD-I260E molecules were classified as free with only few trajectories classifying as static (Figure 6 B), accentuating the absence of interaction with the membrane or other structures. When comparing Halo-MinD to the monomeric mutants G12V and K16A, the number of static tracks was visibly reduced in the mutants. Since both monomers are unable to dimerise, it is less likely that they can form larger clusters, which we observed in a previous study (44). Between both monomeric mutants, K16A displayed a slightly larger number of static trajectories (Figure 6 B), which is in alignment with BLI data, where the K16A mutant dissociates much less effective from the membrane (Figure 3 C & D, Table 2).

Next, we examined the average intracellular mobility of Halo-MinD compared with the mutant variants by analyzing the mean-squared displacement (MSD, Figure 6 C). In MSD analysis, the traveled distances of recorded trajectories are plotted against a given time lag (Δt) as a function of Δt, here one imaging frame (24 ms), where the slope can indicate the speed (58–60). As expected, the I260E mutant displayed the fastest average mobility, since it does not bind the membrane effectively anymore (Figure 6 C). In contrast, the D40A mutant was the slowest recorded protein, indicated by the only mild incline (Figure 6 C). Since we expect most of the proteins’ population to be membrane bound, and furthermore observed larger foci that suggest cluster formation, this was expected. Halo-MinD wild type protein appeared to be more mobile in comparison (Figure 6 C), as it should exist in both monomeric and dimeric conformation and either freely diffusive in the cytosol or interact with the membrane. Last, the two monomeric mutants G12V and K16A displayed extremely similar average speeds that were both faster than the wild type protein (Figure 6 C). Since these MinD variants cannot form dimers, general diffusion should be faster, as there should be no MinD dimers or oligomers, and possibly no recruitment by MinJ.

**Figure 6:**
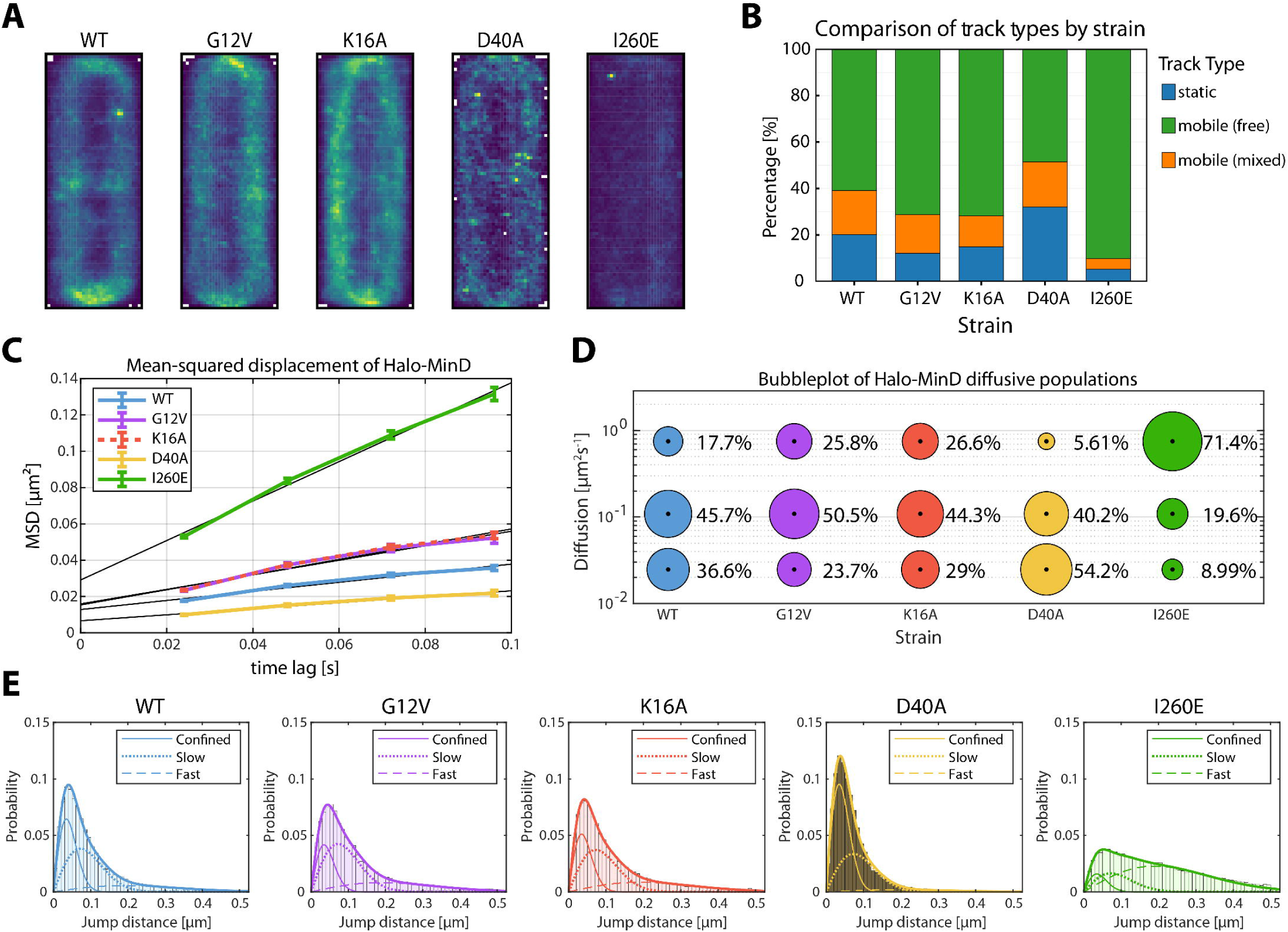
Single-molecule localization microscopy analysis of Halo-MinD and mutants expressed in *B. subtilis*. Exponentially growing *B. subtilis* cells expressing Halo-MinD and variants (n ≥ 48 cells, respectively) were stained with TMR ligand and subsequently imaged. Individual protein trajectories were recorded using SMLM and analyzed with Zen blue (Zeiss), Trackmate, the SMTracker 2.0 software package and manual scripts in R. Minimum track-length 4 frames of 24 ms, with at least 2596 trajectories per strain. (A) Heat map representation of intracellular localization of individual molecules of Halo-MinD and variants, respectively, plotted on normalized cells. Brighter colors indicate higher abundance. (B) Barplot of stationary localization analysis (SLA), comparing different track types within the protein population. Tracks were considered static when not leaving a circular area of 97 nm diameter within 5+ frames. Mobile populations were further divided into free and mixed tracks, where mixed tracks displayed a switch between free and confined movement. (C) Plot of the mean-squared displacement of Halo-MinD and variants over time, fitted with a linear fit excluding the last timelag. (D) Bubble plot displaying single-molecule diffusion rates of the indicated MinD fusions. Populations were determined by fitting the probability distributions of the frame-to-frame displacement (jump distance) data of all respective tracks to a three-component model (fast mobile, slow mobile and confined protein populations). (E) Probability distributions of jump distances of Halo-MinD and variants. Data was fit with a three-component model, indicating confined, slow and fast tracks.

If the underlying movement in MSD analysis can be described by simple Brownian motion, the gradient of the curve is proportional to the diffusion coefficient *D* (60). We know, however, that MinD interacts at least with MinC, MinJ, the membrane and itself, in combination with the fact that bacterial cells exhibit natural confinement. With these information one cannot assume simple Brownian motion, therefore we have chosen jump distance (JD) analysis as a more accurate assessment to describe Halo-MinD dynamics (Figure 6 D & E). In JD analysis, the distances molecules travel between subsequent frames are plotted in a probability distribution (Figure 6 E), and diffusion coefficients are obtained by curve fitting (60), in this case with a three-component model separating fast, slow and confined molecules (Figure 6 D & E). To accurately compare population sizes, the simultaneous option was chosen in the SMTracker software, fixing the diffusion coefficients for every MinD variant (57). To also obtain individual, variant specific diffusion coefficients, the same data was additionally fitted individually (non-simultaneous SQD analysis in SMTracker2, Table 4).

**Table 4:** Diffusion coefficients obtained from non-comparative SQD analysis of Halo-MinD and variants fitted with three populations.

| Strain | Fast diffusive [$\mu\text{m}^2 \text{ s}^{-1}$] | Slow diffusive [$\mu\text{m}^2 \text{ s}^{-1}$] | Confined [$\mu\text{m}^2 \text{ s}^{-1}$] |
| --- | --- | --- | --- |
| **MinD WT** | 0.60 (22.3 %) | 0.084 (50.4 %) | 0.020 (27.3 %) |
| **MinD-G12V** | 0.68 (28.5 %) | 0.10 (47.5 %) | 0.026 (24.0 %) |
| **MinD-K16A** | 0.70 (27.3 %) | 0.11 (43.9 %) | 0.024 (28.8 %) |
| **MinD-D40A** | 0.44 (13.7 %) | 0.063 (50.0 %) | 0.021 (36.3 %) |
| **MinD-I260E** | 0.76 (73.4 %) | 0.095 (20.3 %) | 0.022 (6.3 %) |

Percentages indicate relative population size

Generally, the results of JD analysis correlated well with our *in vitro* data. When dissecting the different populations in the wild type, more than a third of all proteins appear to be confined (36.6%, Figure 6 D), which likely comprises membrane bound protein that is further stabilized by protein-protein interactions via lateral interaction of MinD with itself, and possibly MinC and MinJ. The slow mobile population was the largest (45.7%, Figure 6 D), and likely summarizes different conformations of MinD that interact with the membrane, but are still diffusive and not captured or stabilized through further interactions. The third, fast diffusive population (17.7%, Figure 6 D) likely reflects cytosolic MinD. These results stress the importance of the MinD ATPase cycle for its localization and dynamics, as these are severely altered in the MinD mutants.

In detail, the probability distributions of Halo-MinD-D40A and I260E (Figure 6 E) illustrate the strong shift between MinD molecules in a confined (D40A) and a diffusive state (I260E) best. When examining the different populations, most Halo-MinD-D40A molecules were classified as confined (54.2%) and slow mobile (40.2%), whereas I260E only displayed ∼9% of confined molecules (Figure 6 D), matching prior results and expectations of the membrane binding mutant. When comparing D40A to wild type Halo-MinD, the largest shift is observed between the fast mobile and confined populations, shifting towards a distinctively more confined regiment in the D40A mutant (36.6% WT, 52.2% D40A; Figure 6 D) and visible disappearance of the fast population in the probability distribution (Figure 6 E). These results are in line with the previous observations, as a trapped dimer of MinD will remain at the membrane at much higher efficiency (Figure 5 E & Table 2), and appeared to form foci and clusters (Figure 3 B, Figure 6 A). G12V and K16A mutants on the other hand showed an increase in fast mobile populations (WT 17.7%, G12V 25.8%, K16A 26.6%; Figure 6 D E), which could be attributed to a reduced interaction with MinD interaction partners (MinC, MinD, MinJ and plasma membrane). This notion is supported by the decrease in confined molecules for both G12V and K16A mutants compared to the wild type (WT 36.6%, G12V 23.7%, K16A 29%; Figure 6 D & E), which are expected to be membrane bound and likely stabilized through other protein-protein interaction, which may be reduced or absent in the monomeric conformation. This notion is also supported by the absence of a MinD gradient in epifluorescence images of G12V and K16A mutants (Figure 3 B), a pattern usually formed through sequestering via MinJ. Finally, when comparing both monomeric variants, the K16A mutant displays a noticeable shift between the slow diffusive (G12V 50.5%, K16A 44.3%; Figure 6 D) and confined populations (23.7% G12V, 29% K16A; Figure 6 D). The larger confined population of Halo-MinD-K16A is consistent with the BLI results (Figure 5 D, Table 2), where K16A appears to have a lower dissociation from the membrane, increasing its likelihood of confinement, possibly caused by the inability to bind ATP.

In summary, these results emphasize the importance of the MinD ATPase cycle for functionality of the Min system in *B. subtilis*. Even though *B. subtilis* MinD is not capable of oscillation, its intracellular dynamics are highly dependent on the functionality of the MinD ATPase domain and its catalytic site, which is reflected by the variety of localization and interaction defects displayed by the MinD mutants on a molecular level, that all result in severe division anomalies.

### MinC and MinJ have different effects on MinD protein dynamics in SMLM

MinC and MinJ interact with MinD *in vivo*, suggesting protein dynamics should be affected by this. However, in our ATPase and BLI experiments, MinC had only minor effects on ATPase activity and membrane binding kinetics of MinD, while the PDZ domain slightly enhanced ATPase activity without altering membrane binding kinetics of MinD. To validate these observations *in vivo* and to identify differences between MinD interacting with a cytosolic PDZ-domain or a full-length MinJ, we performed the same SMLM workflow and analysis pipeline described in the previous section on cells expressing Halo-MinD in Δ*minC* and Δ*minJ* backgrounds and compared them to the wild type background (Figure 7, Table 5).

**Figure 7:**
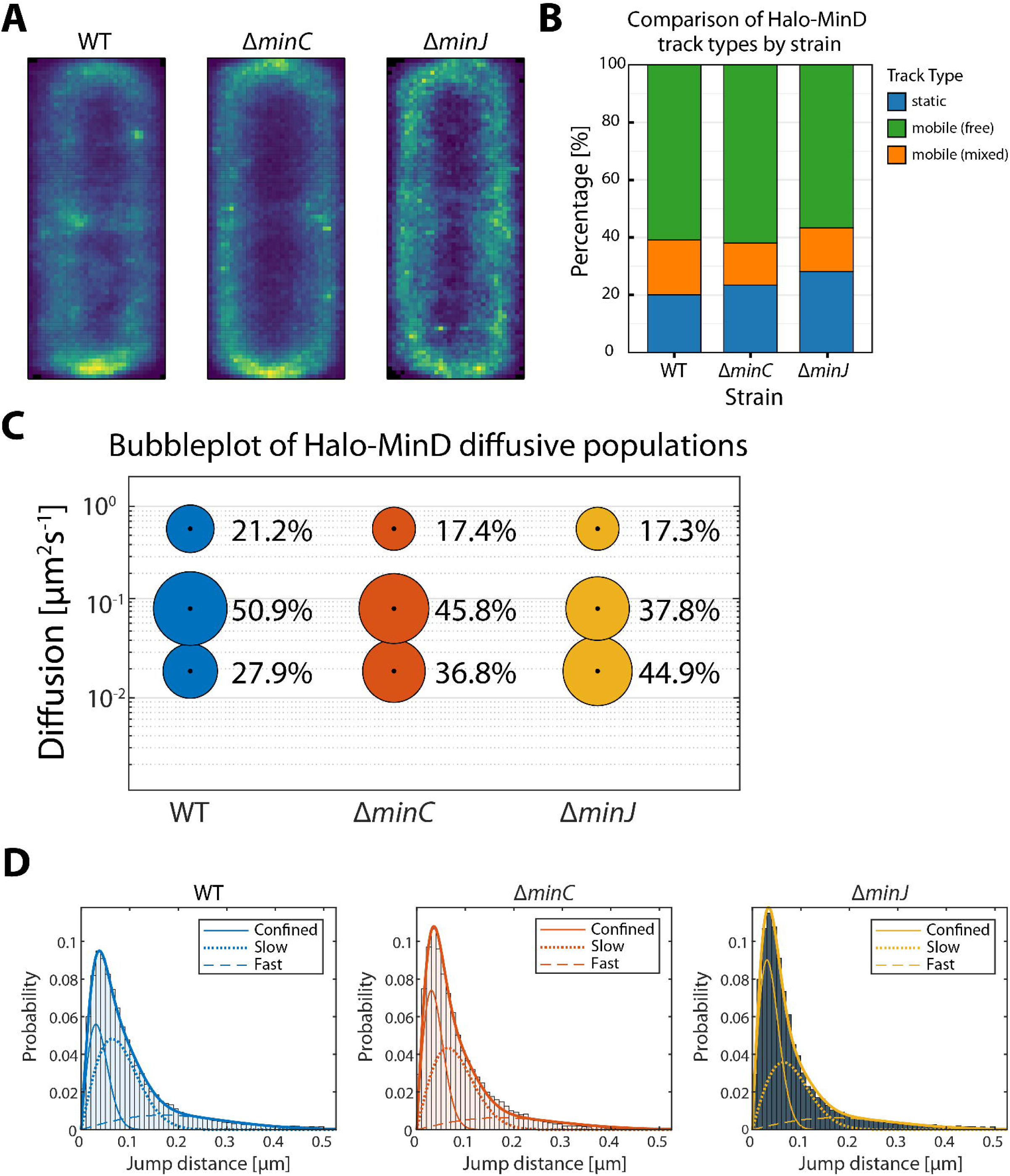
Single-molecule localization microscopy analysis of Halo-MinD expressed in *B. subtilis* wild type and mutant backgrounds. Exponentially growing *B. subtilis* cells expressing Halo-MinD in the indicated genetic backgrounds (n ≥ 115 cells, respectively) were stained with TMR ligand and subsequently imaged. Individual protein trajectories were recorded using SMLM and analyzed with Zen blue (Zeiss), Trackmate, the SMTracker 2.0 software package and manual scripts in R. Minimum track-length 4 frames of 24 ms, with at least 2688 trajectories per strain. (A) Heat map representation of intracellular localization of individual molecules of Halo-MinD in the indicated genetic background, respectively, plotted on normalized cells. Brighter colors indicate higher abundance. (B) Barplot of stationary localization analysis (SLA), comparing different track types within the protein population. Tracks were considered static when not leaving a circular area of 97 nm diameter within 5+ frames. Mobile populations were further divided into free and mixed tracks, where mixed tracks displayed a switch between free and confined movement. (C) Bubble plot displaying single-molecule diffusion rates of Halo-MinD in different genetic backgrounds. Populations were determined by fitting the probability distributions of the frame-to-frame displacement (jump distance) data of all respective tracks to a three components model (fast mobile, slow mobile and confined protein populations). (D) Probability distributions of jump distances of Halo-MinD in different genetic backgrounds. Data was fit with a three-component model, indicating confined, slow and fast tracks.

**Table 5:** Diffusion coefficients obtained from non-comparative SQD analysis of Halo-MinD in different genetic backgrounds fitted with three populations.

| Strain | Fast diffusive [$\mu\text{m}^2 \text{s}^{-1}$] | Slow diffusive [$\mu\text{m}^2 \text{s}^{-1}$] | Confined [$\mu\text{m}^2 \text{s}^{-1}$] |
| --- | --- | --- | --- |
| **WT** | 0.57 (22.1 %) | 0.083 (50.2 %) | 0.019 (27.7 %) |
| ***ΔminC*** | 0.31 (29.3 %) | 0.059 (44.3 %) | 0.01 (26.4 %) |
| ***ΔminJ*** | 0.6 (19 %) | 0.069 (41.2 %) | 0.018 (39.7 %) |

Percentages indicate relative population size

When comparing the average Halo-MinD distribution between the respective strains, the Δ*minC* mutant exhibited a similar intracellular localization pattern as the wild type, with strong membrane enrichment and highest concentrations at midcell and poles (Figure 7 A). However, Δ*minC* cells interestingly exhibited a slightly reduced cytosolic signal and a corresponding increased membrane association. As expected, Δ*minJ* cells lacked the characteristic septal accumulation. Instead, Halo-MinD was distributed relatively uniformly along the membrane, consistent with MinJ being required for efficient sequestering of MinD to the future division site and cell poles.

In the stationary localization analysis (Figure 7 B), Halo-MinD in the Δ*minC* strain displayed only a modest increase in the static fraction compared to wild type (23.4 % from 20.1 %). This shift can be attributed to a reduction in the mixed, likely membrane-interacting population (from 19.0 % to 14.7 %), whereas the freely mobile fraction remained essentially unchanged (61 % to 62 %). These results indicate that in the absence of MinC, less MinD moves along the membrane and a slightly larger proportion becomes immobilized. Although the effect size is small, it suggests that MinD may form larger or slightly more stable membrane-associated assemblies when MinC is absent *in vivo*. In the Δ*minJ* background, the static Halo-MinD population surprisingly increased (20.1 % to 28.2 %), together with respective reductions in both the freely diffusing cytosolic pool (61.0 % to 56.6 %) and the mixed, likely membrane-interacting population (19.0 % to 15.1 %). This redistribution suggests that MinJ contributes to efficient MinD turnover on the membrane, in line with MinD ATPase data with His-PDZ (Figure 7). In the absence of MinJ, MinD appears to remain membrane-bound for longer, consistent with impaired detachment or reduced cycling between membrane-associated states. One possible explanation is that MinD may form more stable or aggregation-prone assemblies when MinJ is absent, whereas the presence of MinJ could promote more dynamic membrane association and hence sharper local accumulations and cycling of MinD, necessary for enriched polar and septal localization. Jump-distance analysis with fixed diffusion coefficients corroborated these findings (Figure 7 C & D), showing that Δ*minJ* cells contained a substantially enlarged confined population of MinD (27.9 % vs. 44.9 %), together with a large reduction of the slow diffusive population (50.9 % vs. 37.8 %). Fitting the data without fixed diffusion coefficients produced the same trend (Table 5), indicating that the shift reflects MinD spending more time in the respective state rather than actual changes in the underlying diffusive coefficients.

In the Δ*minC* strain, Halo-MinD shows a slight redistribution toward the confined state, with an increased fraction of very slow or immobile molecules (27.9 % vs. 36.8 %) and a corresponding decrease in both the fast and intermediate populations (50.9 % vs. 45.8 % and 21.2 % vs. 17.4 %, respectively, Figure 7 C & D). SQD analysis without fixed diffusion coefficient (Table 5) shows that the diffusion coefficients of the fast and intermediate populations are reduced compared with wild-type, indicating that the mobile molecules themselves move more slowly in average. Combined, this could suggest that in the absence of MinC, membrane-bound MinD is more prone to forming clusters or larger aggregates, which restrict mobility and reduce the efficiency of its membrane cycling.

Together, these data suggest that MinJ promotes productive membrane cycling of MinD, and in its absence, MinD accumulates more often in a confined, likely non-productive membrane-associated state, whereas MinC plays a comparatively minor regulatory role, where it passively contributes to MinD mobility by preventing excessive clustering and facilitating proper membrane-associated dynamics.

## Discussion

In bacteria, the oscillation of MinCDE in *E. coli* is one of the best studied examples of a reaction-diffusion system, and the two core components MinD and MinE cycle between membrane and cytosol in an ATP-dependent manner, generating a standing wave with a time-resolved minimum at midcell (61, 62). Here, binding and unbinding of the components to the membrane surface influences the diffusion coefficients and controls the dynamic activity of MinC (63). While the *E. coli* MinCDE system is one of the best-studied examples of intracellular pattern formation in bacteria (62–65), the Min system in *B. subtilis* remains less well characterized. In a previous study (44), we suggested that gradient formation depends on ATPase-driven MinD dynamics, backed by PALM and FRAP analysis as a basis for a mathematical model. However, due to a lack of biochemical data, we had to assume that the *B. subtilis* MinD cycles between membrane-bound and cytosolic localizations based on ATP-binding and hydrolysis, similar to *E. coli* MinD (44). A further gap in our knowledge of the *B. subtilis* Min system concerned the activation mechanism of MinD in absence of a MinE homolog.

To address this gap, we first analyzed *B. subtilis* MinD biochemically *in vitro*. Purified MinD showed basal ATPase activity that drastically increased upon liposome addition. Although *E. coli* MinD similarly requires lipids, it additionally depends on MinE for full activation (47). The reported maximum ATPase rate, around 20 nmol mg⁻¹ min⁻¹ (47, 48), was nearly identical to that of *B. subtilis* MinD, indicating that membrane binding alone is sufficient to trigger robust ATP hydrolysis in the *Bacillus* system. When comparing to other MinD/ParA ATPases that have been biochemically characterized, the activity level of *B subtilis* MinD is on the lower end of the spectrum with a *k*_cat_ of 36.27 h^−1^. Soj, the ParA homologue in *B. subtilis*, reaches more than 2-fold higher catalytic activity (83.6 h^-1^) when fully activated by supplementing Spo0J (ParB) and DNA (66), similarly to Soj in *Thermus thermophilus* combined with Spo0J and *parS* containing DNA (53). Studies investigating the activity levels of ParA in *Caulobacter crescentus*, purified in presence of ATP, determined a *k*_cat_ of 0.79 min^-1^ (47.4 h^-1^) in absence of ParB, increasing almost 5-fold to 3.7 min^-1^ (222 h^-1^) in presence of ParB (67). A more recent study found the *C. crescentus* ParA *k*_cat_ increase from 5.8 hr^−1^ to 120 hr^-1^ upon addition of ParB (68).

These findings suggest that membrane association is the primary trigger for MinD ATPase activity in *B. subtilis*. Specifically, the comparison of maximum velocities, activity stimulation factors, and K_M_ values between the *E. coli* and *B. subtilis* MinD proteins suggests that a functional homolog of the MinE protein is not necessary in *B. subtilis*. At the same time, our biochemical data indicate that MinJ/PDZ increases MinD ATP hydrolysis rates at least to a modest extent, (between 22% and 37%, Figure 2 B), pointing to a broader regulatory influence of MinJ on MinD that becomes evident from additional mechanistic analyses discussed below. In contrast, the presence of MinC led only to a slight decrease in MinD ATPase activity in our APTase assays (∼ 5%, Figure 2 A). This modest inhibitory effect suggests that MinC does neither directly stimulate MinD turnover nor inhibit it, and may instead stabilize MinD in a conformation or assembly state with mildly reduced catalytic activity, also discussed below.

With the goal of testing binding of isolated MinD to the membrane, we designed a Bio-layer interferometry experiment in which we immobilized liposomes and measured binding of purified MinD proteins. Importantly, without addition of ATP (and ADP present from purification) we observed only very weak MinD binding to the liposomes (Figure 5–figure supplement 2), indicating that MinD membrane binding and dissociation is coupled to the ATP hydrolysis cycle or the nucleotide binding state. In presence of ATP, MinD readily binds to the liposomes with high affinity. The ParA/MinD family is well studied, and mutations producing strictly monomeric (G12V, K16A), dimeric (D40A), or membrane-binding-deficient (I260E) MinD variants are well established (51–53). The most intriguing observation of the binding experiments is that MinD monomers can apparently bind to the liposome membrane *in vitro*. Both monomeric mutants (G12V and K16A) are readily binding to the membrane, but the release of the proteins from the membrane surface differs drastically. While the K16A mutant, unable to bind ATP, has a slow membrane release rate (*k_off_* rate of only 2.77*10^-3^ s^-1^), the G12V monomeric mutant, which can still bind ATP, is released fast from the membrane (*k_off_* rate 47.8*10^-3^ s^-1^). The D40A mutant of MinD, which is thought to be a stable ATP-bound dimer, releases with very slow kinetics (*k_off_* rate 1.76*10^-3^ s^-1^). This indicates that ATP hydrolysis accelerates MinD release from the membrane and that ATP binding itself influences MinD association and dissociation kinetics, although we cannot entirely exclude that the mutations introduce subtle structural changes that affect membrane affinity. These findings crucially support a reaction diffusion model in which the MinD ATPase cycle is essential for correct pattern formation of *B. subtilis* MinD. When testing the impact of MinC on MinD membrane-binding kinetics, no significant differences were observed, suggesting that MinC does not substantially influence the binding dynamics of MinD, in-line with the collective view of MinC as a passenger rather than a driver of MinD dynamics (63). It would have been desirable to study the interaction of MinD with reconstituted full-length MinJ proteins to specifically investigate the influence that MinJ has on membrane interaction of MinD. Unfortunately, we did not succeed in the purification of a full length MinJ protein so far. MinJ contains seven predicted transmembrane helices and displays a pronounced instability, rendering purification difficult. While the binding kinetics were unaffected during BLI, our *in vitro* ATPase experiments combining the soluble PDZ domain of MinJ with MinD already demonstrate that MinJ possibly modulates MinD:membrane interactions to some extent through an effect on the hydrolysis efficiency.

To clarify this point and to better understand the isolated MinD membrane interaction under *in vivo* conditions, we turned to state-of-the-art microscopy (SMLM) (69, 70).

SMLM data comparing the behavior of MinD in wild type vs. Δ*minJ* background (Figure 7) collectively suggest that MinJ stabilizes correct pattern formation of MinD. By promoting efficient membrane detachment and maintaining MinD in a dynamically cycling state, MinJ appears to ensure that MinD remains properly redistributed throughout the cell cycle rather than becoming sequestered in non-productive clusters. This organizational role likely underlies the ability of MinJ to position MinD, and thus MinC, at the appropriate cellular sites (38, 39), thereby supporting robust spatial regulation of division. However, a short-lived MinJ mediated stabilization of MinD-dimers at the membrane would be comparable to the stabilizing effect that MinE has on MinD oligomers in *E. coli* (71, 72), and was previously observed in our *in vivo* FRAP studies (44). In contrast to MinJ, MinC did not seem to have a major impact on MinD dynamics *in vivo*. When observing MinD in absence of *minC*, the main effect was a subtle reduction in MinD cycling activity, shifting a small fraction of MinD into more static or confined membrane-associated states. In the wild type situation, MinC may passively restrict uncontrolled aggregation by capping or destabilizing certain interfaces, or conversely transiently stabilize short-lived assemblies required for regulation.

The locked MinD dimers (D40A mutant) frequently nucleated into larger membrane clusters. This cluster formation is underlined by the drastic increase of the confined/immobile population (54.2%) of D40A in SMLM. This increase in the static population comes to the expense of freely diffusive protein (∼5.6 %), indicating that the vast majority of MinD D40A is membrane associated. We observed a clear enrichment of these MinD clusters at the cell poles and septa, lending support to the notion that MinD dimers are specifically stabilized/recruited by MinJ. This interpretation is supported by previous data showing that in absence of MinJ, MinD proteins are randomly localized along the membrane in clusters, but not enriched at poles or (in a *minJ* mutant background rare) septa (40). In line with this idea, the monomeric MinD variants (G12V, K16A) are evenly bound to the membrane surface along the entire cell length.

The fact that most monomeric MinD appears membrane associated *in vivo* also supports our *in vitro* measurements in which monomeric MinD variants efficiently bind the membrane, a notion strongly supported by a recent report that examined localization of the same monomeric variants in epifluorescence microscopy (73). The time-resolved MinD gradient is lost in all mutants - including D40A - which should still be recruited by MinJ, suggesting the ATPase cycle and thus MinD cycling is indeed necessary for proper formation of a gradient, as suggested before (44). Our data are in perfect agreement with recent *in vitro* data collected with the *E. coli* MinD using mass-sensitive particle tracking (MSPT) (56). MSPT analysis revealed that *E. coli* MinD forms lateral interactions, leading to higher order oligomers (trimer, tetramer and even higher oligomers). It was speculated that dimerization of *E. coli* MinD at the membrane leads to conformational changes opening an additional interaction surface of MinD for lateral interaction (56). These lateral MinD interactions were thought to recruit further monomeric MinD from solution to the membrane, thereby forming a nucleation for larger clusters. The MinD cluster formation that we observed in *B. subtilis* using SMLM (44) are likely *in vivo* evidence for this cooperative membrane binding. It should be noted that the MSPT analysis was not sensitive enough to detect monomer binding to the membrane, but this was explicitly not ruled out (56). Addition of a second amphipathic helix to the *E. coli* MinD however, was shown to promote monomer binding of *E. coli* MinD to the membrane (49). Thus, the difference between the *E. coli* and *B. subtilis* MinD is likely the higher membrane binding capacity of the *Bacillus* protein. The observed cooperativity in membrane binding and thus nucleation, as shown for the *E. coli* MinD in vitro (49, 56) and here for the *B. subtilis* MinD *in vitro* and *in vivo*, is required for pattern formation by conferring nonlinearity (44, 74, 75).

Consistent with this model, our SMT data revealed three major dynamic states of MinD: fast cytosolic diffusion, a slower membrane-associated pool, and a confined fraction corresponding to clustered or multimeric assemblies. Most of the membrane-binding-deficient mutant I260A shifted drastically into the fast, cytosolic population, validating this assignment. Conversely, the locked-dimer (D40A) mutant showed a pronounced increase in confined, slow-moving molecules, in line with its strong tendency to nucleate clusters. Monomeric variants (G12V, K16A) displayed reduced confinement, supporting the view that dimerization is required for higher-order assembly or recruitment. Together, these observations indicate that MinD dynamically cycles between cytosol and membrane in an ATP-dependent manner. Thus, we conclude that membrane bound dimers of MinD have a much stronger tendency to interact with and recruit binding partners (including self-nucleation). This finding is further supported by a recent report that analyzed MinD dynamics *in vivo* using diffraction limited microscopy (73). In this study, a MinD D40A mutant hyper-recruited MinC, leading to cluster formation of MinCD at the membrane. Deletion of MinC restored the MinD gradient formation, indicating that MinC might indeed be part of a MinC:MinD co-polymer, as suggested for the *E. coli* system (76). This assumption is in good agreement with our *in vivo* findings, where self-nucleation seems to increase drastically for the D40A mutant or in absence of MinJ. In contrast, monomeric G12V and K16A mutants do not display this behavior, in line with monomeric MinD variants not being able to interact with MinC, which was similarly observed through widefield microscopy in the same study (73).

Collectively, our data reveal that the MinD ATPase cycle is essential for regulated membrane binding and release, which is crucial for the formation of a sharp MinD gradient at the septum (44). However, in contrast to *E. coli* MinD, monomeric *B. subtilis* MinD can bind the membrane with higher affinity, suggesting that MinD also dimerizes at the membrane surface. Dimer formation is essential for ATPase activity, as indicated by our *in vitro* measurements, and is necessary for efficient membrane detachment. The main characteristics of the *E. coli* MinD with respect to membrane binding, self-interaction leading to nucleation, and membrane detachment are therefore shared in the *B. subtilis* protein. The difference, that ATP hydrolysis of the *B*. *subtilis* MinD does not require a trigger other than membrane binding, makes sense in a system that does not need to oscillate, but rather accumulate and sustain a sharp gradient at the current site of division. However, the balance of a dynamic membrane cycle of MinD seems to be fine-tuned by the interaction with MinC and especially MinJ, which appears to keep MinD in a productive state and protects it from aberrant aggregation, while recruiting it towards poles and septum, the intended site of action of MinC.

While these findings elucidate the functional principles of the *B. subtilis* Min system further, it is not clear if and how broadly they can be inferred on other rod-shaped, Gram-positive bacteria that possess a Min system. While Min systems are often summarized into either oscillating MinCDE systems or DivIVA dependent gradients, the group of spore-forming Firmicutes appear to have rather individual sets of Min proteins. Often, they contain homologs from both systems, presumably necessary to individually regulate and allow the switch between symmetric (vegetative) and asymmetric (sporulation) division (77). A recent study demonstrated e.g. that *Clostridioides difficile* MinDE proteins expressed in *B. subtilis* can display oscillatory behavior, interfering with sporulation (78), while DivIVA in *C. difficile* appears to directly interact with MinD without a MinJ-like bridge protein (79). Furthermore, a very recent study identified another protein (MrpA) in *C. difficile* that directly interacts with MinD, an interaction leading to aberrant cell division as well as affecting colony morphology (80). The extent to which these mechanisms operate simultaneously or are differentially utilized under specific physiological conditions, such as vegetative growth or sporulation, remains an open question. Our findings on the Min system in *B. subtilis* therefore likely represent only one end of a broad spectrum of bacterial division site selection mechanisms.

## Materials and Methods

### Key Resources Table

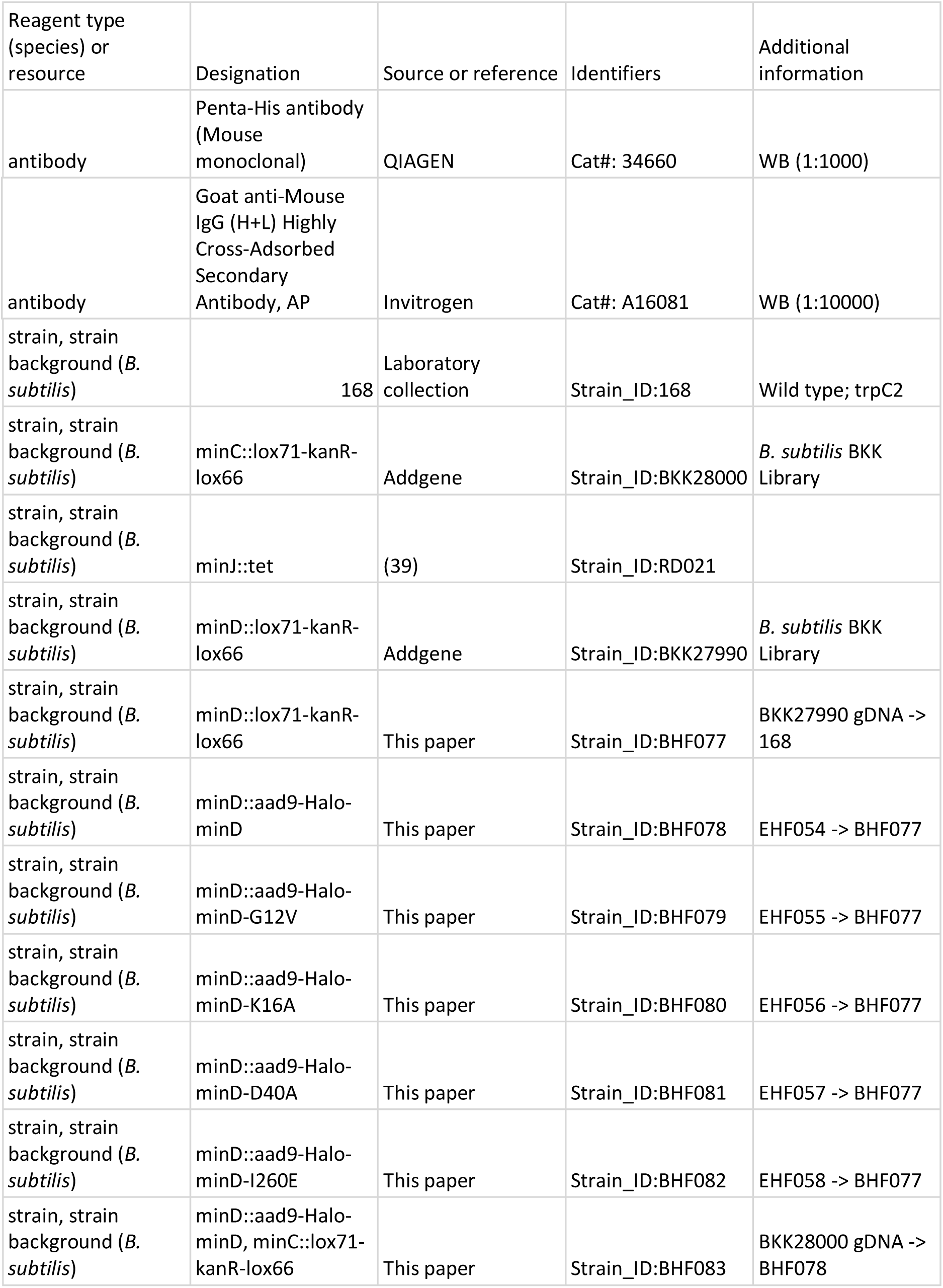
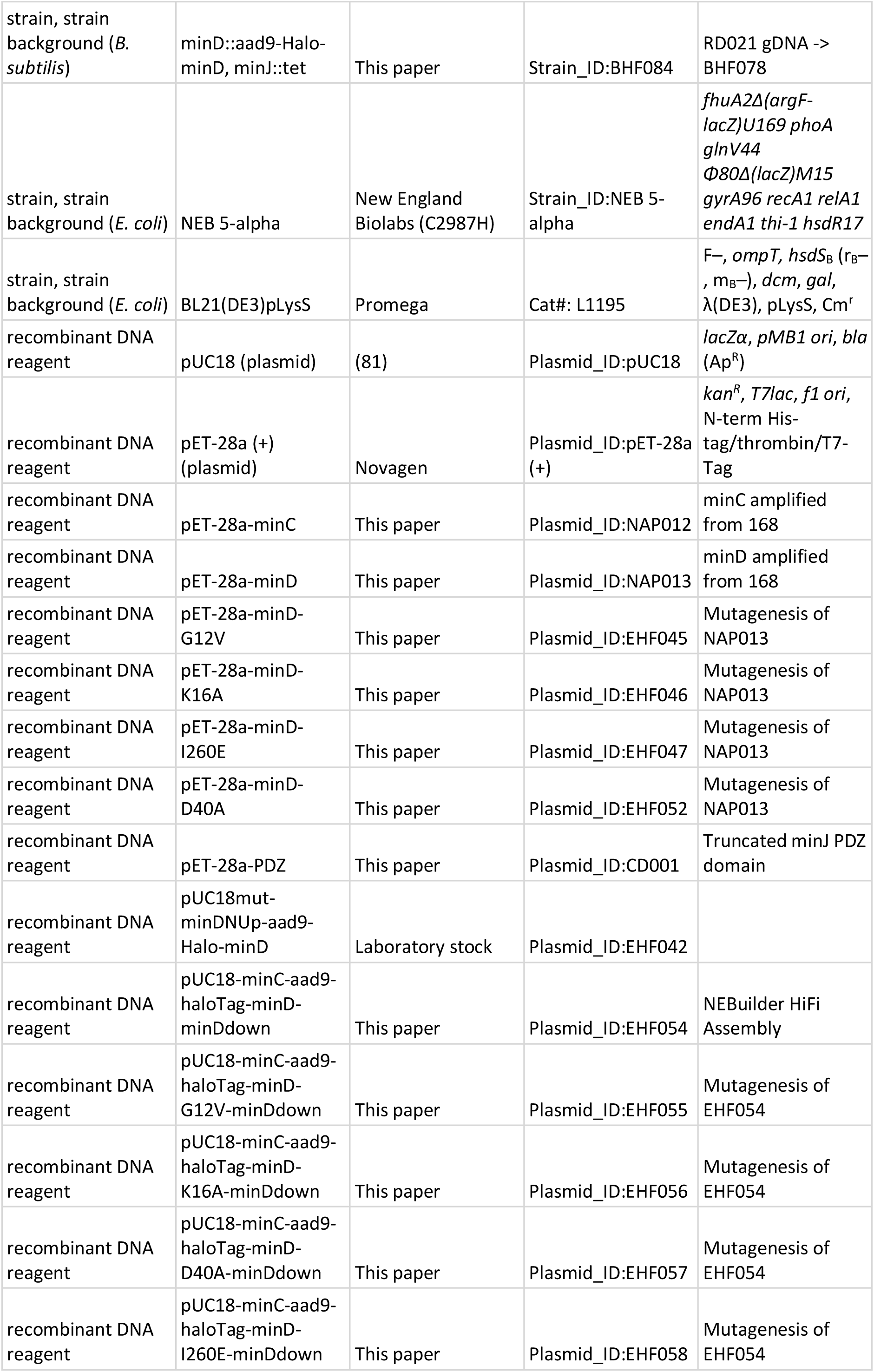
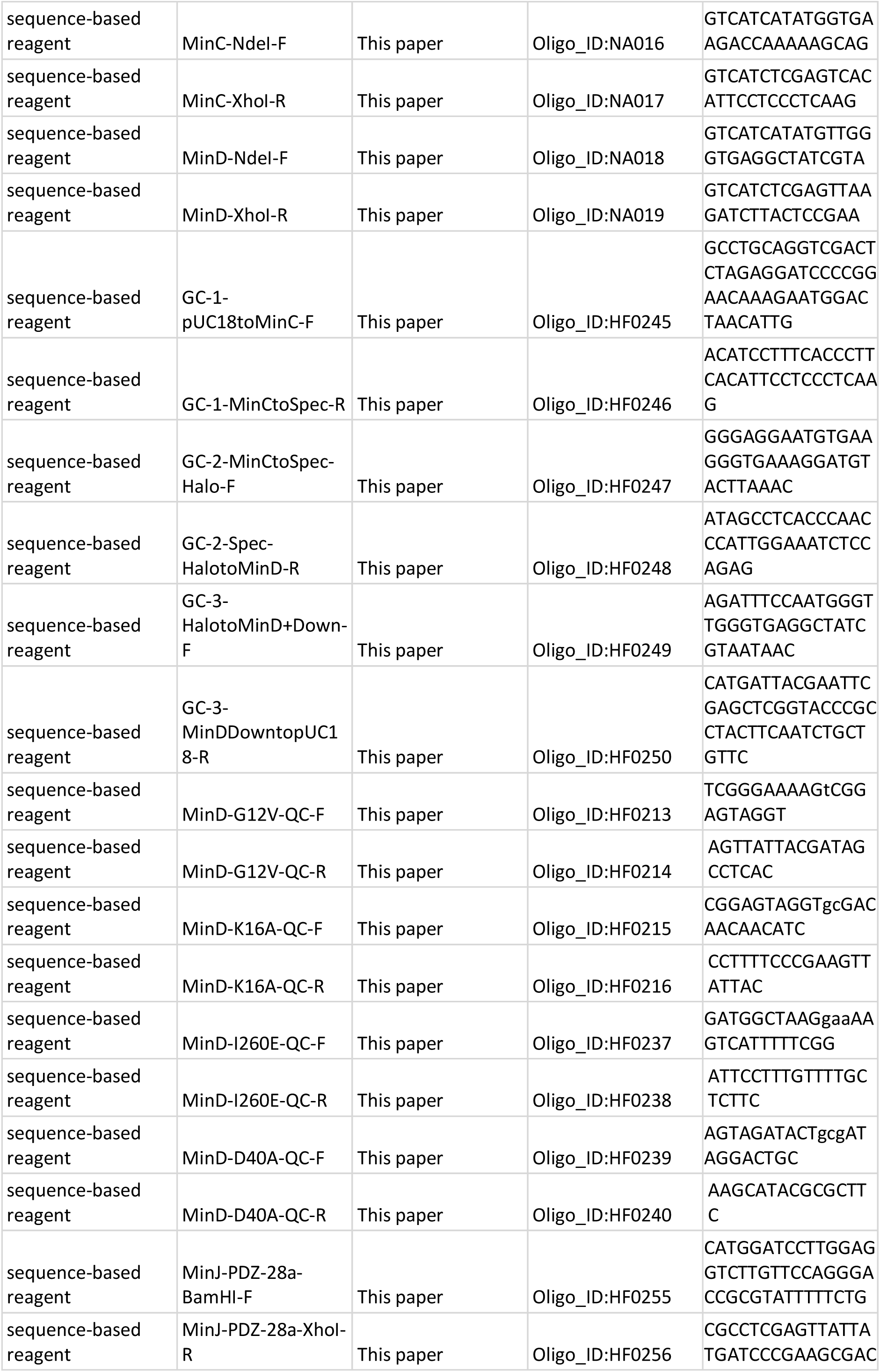
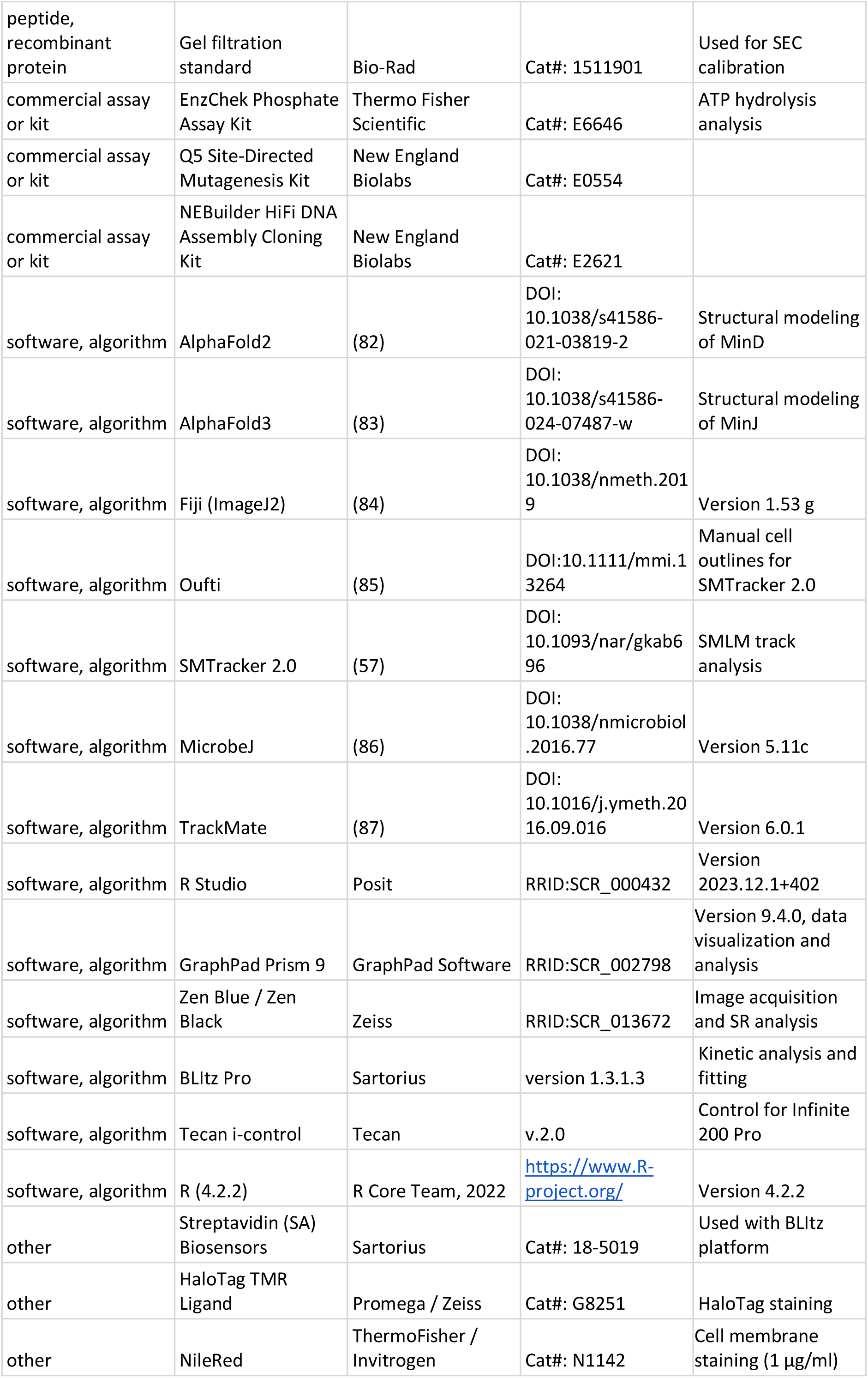
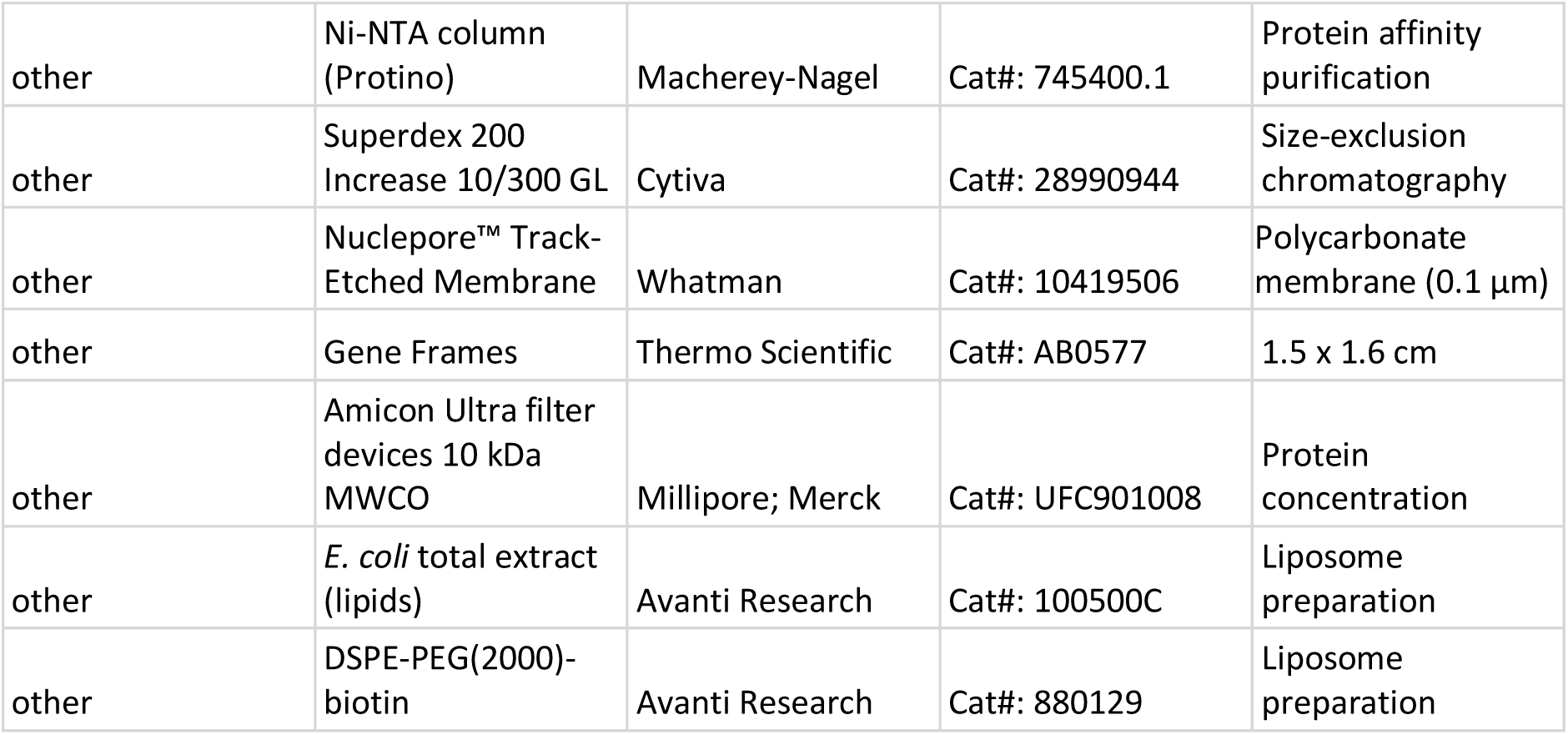

### Bacterial strains, plasmids and oligonucleotides

Plasmid and strain construction of all relevant constructs is described in the Appendix. Oligonucleotides, plasmids and strains used in this study are listed in Supplementary table 1, Supplementary table 2 and Supplementary table 3, respectively, as well as in the key resources table.

### Media and growth conditions

*E. coli* was grown on lysogeny broth (LB) agar plates containing 10 g L^-1^ tryptone, 5 g L^-1^ yeast extract, 10 g L^-1^ NaCl and 1.5% (w/v) agar at 37°C overnight using appropriate selection (kanamycin 50 µg ml^-1^, chloramphenicol 35 µg ml^-1^).

*B. subtilis* was grown on nutrient agar plates using commercial nutrient broth and 1.5% (w/v) agar at 37°C overnight. To reduce inhibitory effects, antibiotics were only used for transformations and when indicated, since allelic replacement is stable after integration (kanamycin 5 µg ml^-1^, spectinomycin 100 µg ml^-1^).

For microscopy, *B. subtilis* was inoculated to an OD_600_ 0.05 from a fresh overnight culture and grown in MD medium - a modified version of Spizizen Minimal Medium (88) – at 37°C with aeration in baffled shaking flasks (200 rpm) to OD_600_ 0.5-0.8. MD medium contains 10.7 mg ml^−1^ K_2_HPO_4_, 6 mg ml^−1^ KH_2_PO_4_, 1 mg ml^−1^ Na_3_ citrate, 20 mg ml^−1^ glucose, 50 µg ml^−1^ L-tryptophan, 11 µg ml^−1^ ferric ammonium citrate, 2.5 mg ml^−1^ L-aspartate and 0.36 mg ml^−1^ MgSO_4_ and was always supplemented with 1 mg ml^−1^ casamino acids. Subsequently, cultures were diluted to OD_600_ 0.05 in fresh, pre-warmed MD medium and grown to OD_600_ 0.5 (exponential phase).

### Purification of His-MinC, His-PDZ, His-MinD and variants

For heterologous expression of His-MinC, His-PDZ and His-MinD and its variants, freshly transformed BL21(DE3) cells carrying the respective pET-28a expression vector (Supplementary table 2) as well as pLysS (His-MinC, His MinD and variants only) were transferred to LB medium with appropriate selection (kanamycin 50 µg ml^-1^; chloramphenicol 35 µg ml^-1^ in presence of pLysS) and grown overnight (37°C, 200 rpm). The next day, fresh LB medium was inoculated 1:150 from the overnight culture and grown to an OD₆₀₀ of 0.5 at 37 °C and 200 rpm. Since His-PDZ tended to form inclusion bodies under these conditions, its expression culture was instead grown at 18 °C and 120 rpm without IPTG induction. For all other constructs, protein expression was induced with 1 mM IPTG. Cells expressing His-MinC and all His-MinD variants were harvested after 3 h of induction, whereas His-PDZ cultures were collected after overnight incubation. Cells were harvested by centrifugation (4 °C, 5000 × g, 15 min) and stored at -80°C. For purification, a protocol of *E. coli* MinD purification was adapted from (46): the cell pellet was resuspended on ice in lysis buffer (50 mM NaH_2_PO_4_, 500 mM NaCl and 10 mM imidazole; pH 8.0), supplemented with protease inhibitor cocktail (Roche), 0.2 mM Mg^2+^-ADP and 0.4 mM TCEP (His-MinD and variants only) and ∼250 U mL^-1^ DNase I (Roche), and subsequently lysed in a pre-cooled (4°C) French pressure cell (Amico; 3 x 1,200 psi). The debris was then removed via centrifugation (10,000 g, 30 min, 4°C) and the supernatant was passed through a 1 ml Ni-NTA column (Protino, Macherey-Nagel) utilizing liquid chromatography (ÄKTApurifier). Here, the resin was washed with 10 ml binding buffer A (50 mM NaH_2_PO_4_, 500 mM NaCl, 20 mM imidazol; pH 8) before elution buffer B (50 mM NaH_2_PO_4_, 500 mM NaCl, 250 mM imidazol; pH 8, (and 0.2 mM Mg^2+^-ADP and 0.4 mM TCEP for His-MinD and variants)) was added sequentially at concentrations of 5% (10 ml) 10% (20 ml), and finally 100% (10 ml). Recombinant protein eluate was collected in 0.5 ml fractions. Fractions containing high concentrations of protein (determined by chromatogram analysis, sodium dodecyl-sulfate polyacrylamide gel electrophoresis (SDS-PAGE) and Western blotting using Penta-His antibody (QIAGEN)) were pooled and concentrated using Amicon filter devices with 10 kDa molecular weight cut-off (Millipore; Merck). Additionally, protein aggregates were removed by pelleting them through centrifugation (10,000 g, 10 min, 4°C).

Samples were further purified by size-exclusion chromatography through a Superdex200 Increase 10/300 GL column (Cytiva) at a flow rate of 0.75 ml min^-1^ using gel filtration buffer C (50 mM HEPES-KOH, 150 mM KCl, 10% (v/v) glycerol, 0.1 mM EDTA, pH 7.2). For His-MinD and its variants, buffer C was supplemented with 0.4 mM TCEP and 0.2 mM Mg²⁺-ADP immediately before use. His-PDZ was not subjected to size-exclusion chromatography because it could not be recovered after SEC. Instead, it was used directly following affinity purification. To calibrate the column and allow identification of relevant fractions, a gel filtration standard (Bio-Rad) was run according to the manufacturers protocol and analyzed using the same conditions, respectively. Fractions containing recombinant protein were pooled and either directly used for the respective assays, stored at 4°C for a maximum of one week or flash-frozen and stored at -80 °C. Final protein concentration was determined using Bradford assay with bovine serum albumin (BSA) as standards. Protein purity and stability was confirmed via SDS-PAGE using 12% polyacrylamide gels and Western blotting using a Penta-His antibody (QIAGEN) (Figure 1–figure supplement 1, Figure 2–figure supplement 1, Figure 2–figure supplement 2, Figure 4–figure supplement 1, Figure 4–figure supplement 2, Figure 4–figure supplement 3 and Figure 4–figure supplement 4).

### ATP hydrolysis assays

ATP hydrolysis was analyzed in continuous reactions using the EnzChek Phosphate Assay Kit (Thermo Fisher Scientific), according to the manufacturer’s manual. Reaction volumes were reduced to 100 µl and assayed in flatbottom 96-well plates (Greiner-UV-Star 96-well plates). Reactions were measured at 37 °C continuously every minute over a time-course of 3 h in a Tecan Infinite 200 Pro with the Software Tecan i-control v.2.0. If not indicated differently, the reactions contained either 0-16 µM His-MinD or the respective variant/protein combination as well as 2 mM Mg^2+^-ATP and 0.2 mg ml^-1^ liposomes. Liposomes were made from *E. coli* total extract (Avanti) by first drying and then resuspending and rehydrating lipids in gel filtration buffer C (50 mM HEPES-KOH, 150 mM KCl, 10% (v/v) glycerol, 0.1 mM EDTA). Next, the lipids were extruded 40 times through 100 nm polycarbonate membranes (Nuclepore™ Track-Etched Membrane 0.1 µm, Whatman). Before starting the reactions with 2 mM Mg^2+^-ATP, the plate was preincubated for 10 min in order to eliminate phosphate contamination.

Data were first analyzed with Excel (normalized, subtraction of ATP auto hydrolysis and subtraction of no substrate control), a linear regression was used to determine the hydrolysis rate per minute. Data visualization and further analysis was performed using GraphPad Prism version 9.4.0 for Windows, (GraphPad Software). Statistical comparisons between His-MinD alone (n = 2) and His-MinC/His-PDZ conditions (n = 3 each) were performed using Welch’s t-test (two-tailed), which is appropriate for unequal variances and unequal sample sizes.

### AlphaFold prediction and structure-based analysis

A structural model of *B. subtilis* MinD was calculated using AlphaFold2 (82) installed on a local computer (Figure 3 A), where the structural homology search was performed against the full AlphaFold database as of 11.02.2024 (82).

A structural model of *B. subtilis* MinJ was calculated using AlphaFold3 (83), where the structural homology search was performed against the full AlphaFold database as of 01.02.2025 (83).

### Bio-layer Interferometry

Measurements were performed on the BLItz platform (Sartorius) using Streptavidin (SA) Biosensors (Sartorius, 18-5019). The BLItz device was operated with the Software BLItz Pro (version 1.3.1.3) using the advanced kinetics protocol modified as shown in Table S5 and essentially as described before (89–91).

Beforehand, Liposomes were made from a 33:1 mixture of *E. coli* total extract (Avanti) and DSPE-PEG(_2000_)-biotin (Avanti) by first drying and then resuspending and rehydrating lipids in gel filtration buffer C (50 mM HEPES-KOH, 150 mM KCl, 10% (v/v) glycerol, 0.1 mM EDTA). Next, the lipids were extruded 40 times through 100 nm polycarbonate membranes (Nuclepore™ Track-Etched Membrane 0.1 µm, Whatman). Biosensors were hydrated for at least 10 min in a 96-well plate in 200 µl gel filtration buffer C.

As a starting point, unspecific binding was tested. For this, non-biotinylated His-MinD was used to be loaded onto the Streptavidin biosensor. For the binding assays, all measurements were performed in black reaction tubes with 200 µl gel filtration buffer C or the indicated solution (Supplementary table 4). After measuring the initial baseline, biotinylated liposomes were allowed to bind the biosensor. Another baseline was measured in the same tube as before. For association, 200 µl solutions of His-MinD and variants in different concentrations and/or in combination with His-MinC or His-PDZ, respectively, were exposed to the sensor, first without and later with addition of 2 mM Mg-^2+^ATP (final concentration) just before use. Furthermore, unspecific binding to liposome-saturated sensors was tested using bovine serum albumin (BSA, 16 µM). For dissociation, the sensor was exposed to buffer again. Each measurement was performed with a fresh biosensor.

For BLI experiments involving His-MinD and one of its direct interactors, we first assessed whether His-MinC or His-PDZ can directly interact with His-MinD *in vitro*. To this end, a liposome-coated BLI sensor was first dipped into a 8 µM His-MinD solution. Subsequently, the sensor was exposed to solutions containing either 8 µM His-MinD together with 8 µM His-MinC or 8 µM His-PDZ, respectively. As a negative control, a sample containing 8 µM His-MinD and 8 µM BSA was used. Both His-MinC and His-PDZ induced a significant increase in the equilibrium binding signal upon interaction with immobilized His-MinD, consistent with specific complex formation. In contrast, the BSA control showed only a minimal signal increase, confirming the specificity of the observed binding. To further validate the sensor specificity, individual proteins were tested. Only His-MinD exhibited robust binding to the liposome-coated sensor, while His-MinC and His-PDZ showed negligible signal, indicating non-specific interactions (Figure 5–figure supplement 1). These initial tests support the expected specificity of the MinD:MinC and MinD:PDZ interactions.

Kinetic analysis and fitting were performed by the BLItz Pro Software (version 1.3.1.3). Data was exported and visualized using GraphPad Prism version 9.4.0 for Windows, (GraphPad Software).

The BLItz Pro Software analysis follows a 1:1 binding model, where:

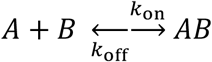

with *k*_on_ = rate of association or “on-rate” and *k*_off_ = rate of dissociation or “off-rate”. Formula for fitting the association (a = slope):

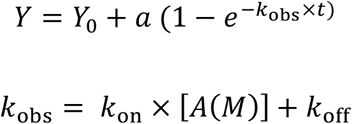

Formula for fitting the dissociation:

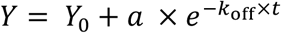

Resulting in the equilibrium dissociation constant *K*_D:_

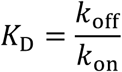

### Fluorescence microscopy

#### Microscopy slide preparation

For epifluorescence imaging, *B. subtilis* cells were mounted on 1.5% MD agarose pads using 1.5 x 1.6 cm “Gene Frames” (Thermo Scientific), and incubated 10 min at 37°C before microscopic analysis.

For SMLM, slides and coverslips were first cleaned by overnight storage in 1 M KOH, carefully rinsed with ddH2O, and subsequently dried with pressurized air. Next, 1.5% (w/v) low melting agarose (Sigma-Aldrich) was dissolved in MD medium. Medium was sterile filtered (0.2 µm pore size) shortly before being used to remove particles. To produce flat, uniform, and reproducible agarose pads, 1.5 x 1.6 cm “Gene Frames” (Thermo Fisher) were utilized, and pads were allowed to solidify for 1 h at room temperature to be used within the next 3 h.

### Staining of HaloTag

For staining, 1 mL of bacterial culture was moved to a 2 mL microcentrifuge tube and incubated with 5 nM (SMLM) or 50 nM (epifluorescence) HaloTag TMR ligand at 37 *◦*C for 10 min. Cells were harvested (4000 g, 3 min, 37 °C) and washed four times in pre-warmed MD medium, before being mounted on the previously described microscopy slides.

#### Epifluorescence imaging

For strain characterization (Figure 3 B), microscopy images were taken on a Axio Observer 7 microscope (Zeiss) equipped with a Colibri 7 LED light source (Zeiss) and an OrcaR^2^ camera (Hamamatsu) using a Plan-Apochromat 100×/1.4 oil Ph3 objective (Zeiss). TMR fluorescence was visualized with a 90 HE LED filter set (QBP 425/30 + 514/30 + 592/30 + 709/100) (Zeiss) and excited at 555 nm (100% intensity of Colibri 7 LED) for 400 ms. The microscope was equipped with an environmental chamber set to 37 °C. Digital images were acquired with Zen Blue (Zeiss) and analyzed and edited with Fiji (84) ImageJ2 (92). For measurement of cell length and minicells, cells were stained with NileRed (1 µg/ml final concentration) at OD_600_ 0.5 and incubated for further 20 minutes (200 rpm, 37°C) before mounting them on an agarose pad. To minimize bias during analysis, cell lengths and number of minicells were identified using the Fiji (84) plugin MicrobeJ 5.11c (86) on phase contrast images with manual correction by screening the results against the simultaneously acquired membrane stain images and sorting out mislabeled cells. For cell length measurements, MicrobeJ modules *options*, *exclude on edges*, *shape descriptor*, *segmentation* and *chain* were used with default settings, except for following parameters in *options* (*area* 1-max; *length* 1-max; *width* 0.6-1; *sinuosity* 1-max; *angularity* 0-0.5) and *chain* (*tolerance* 0.1; *area* 1.5-max). To identify minicells, MicrobeJ modules *options*, *exclude on edges*, *shape descriptor* and *segmentation* were used with default settings, except for following parameters in *options* (*area* 0.15-1; *length* 0.5-1.2; *width* 0.3-1; *circularity* 0.95-max).

### SMLM imaging

SMLM imaging was performed with an Elyra 7 (Zeiss) inverted microscope equipped with two pco.edge sCMOS 4.2 CL HS cameras (PCO AG), connected through a DuoLink (Zeiss), only one of which was used in this study. Cells were observed through an alpha Plan-Apochromat 63x/1.46 Oil Korr M27 Var2 objective, yielding a final pixel size of 97 nm. During image acquisition, the focus was maintained with the help of a Definite Focus.2 system (Zeiss). Fluorescence was excited with a 561 nm (100 mW) laser, and signals were observed through a multiple beam splitter (405/488/561/641 nm) and laser block filters (405/488/561/641 nm) followed by a Duolink SR QUAD (Zeiss) filter module (secondary beam splitter: LP 560, emission filters: EF BP420-480 + BP495-550).

Cells were illuminated with the 561 nm laser (50% intensity) in TIRF mode (62° angle). For each time lapse series 10,000 frames were taken with 20 ms exposure time (∼24 ms with transfer time included) and 50% 561 nm intensity laser.

### SMLM analysis

For average localization analysis (Figure 6 A), a single-molecule localization table was constructed from a 2D Gaussian fitting in the ZEN 3.0 SR (black) software (Zeiss) with a peak mask size of 9 and a peak intensity-to-noise ratio of 6, and further allowing molecules to overlap. Filtering was used to minimize noise, background, and out-of-focus emitters, discarding events that did not emit within a Point spread function size (PSF) of 100-200 nm and did not display a photon count of 50-600. Further analysis was performed using individual Fiji (84) scripts for the cell outlines and R (4.2.2) scripts in R Studio (2023.12.1+402) for the corresponding localization events (93, 94). Here, the remaining events were transformed to align the orientation of all cells in which the signals were detected, to be able to summarize all recorded localizations in one normalized cell per strain. The localization frequency is displayed as a heatmap using the gradual viridis scaling from the ggplot2 R package (95), where the highest frequency (normalized frequency 1) is displayed in yellow and the lowest frequency (normalized frequency 0) in blue (Figure 6 A). The script used to create these heatmaps from SMLM tables was previously published in (96).

For single-molecule tracking, spots were identified with the LoG Detector of TrackMate v6.0.1 (87) implemented in Fiji 1.53 g (84), an estimated diameter of 0.5 μm, and median filter and sub-pixel localization activated. The signal-to-noise threshold for the identification of the spots was set at 7. To limit the detection of particles to single-molecules, frames belonging to the bleaching phase of TMR were removed individually from the time lapses prior to the identification of spots. Spots were merged into tracks via the “Simple LAP Tracker” of TrackMate, with a maximum linking distance of 500 nm and no frame gaps allowed. Only tracks with a minimum length of 5 frames were used for further analysis, yielding a minimum number of total tracks per sample of 2596.

To identify differences in protein mobility and/or behavior, the resulting tracks were subjected to stationary localization analysis (SLA), mean-squared-displacement (MSD) and square displacement (SQD) analysis in SMTracker 2.0 (57), which first required determining cell meshes manually using Oufti (85). SLA analysis was performed with a confinement circle radius of 97 nm and a minimum number of 5 steps to be considered static. The average MSD was calculated for four separate time points per strain (exposure of 20 ms - τ = 24, 48, 72 and 96 ms), followed by fitting of the data to a linear equation. The last time point of each track was excluded to avoid track-ending-related artifacts. The cumulative probability distribution of the square displacements (SQD) was used to estimate the diffusion constants and relative fractions of up to three diffusive states. Diffusion constants were determined simultaneously for the compared conditions, therefore allowing for a more direct population fractions comparison. Additionally, the analysis was performed for each strain non-comparatively, yielding individual diffusion coefficients, displayed in Table 4.

## Acknowledgments

This research was supported by the Deutsche Forschungsgemeinschaft (DFG) within the Transregio Collaborative Research Center (TRR 174) “Spatiotemporal Dynamics of Bacterial Cells”. We thank Leendert Hamoen (Amsterdam) and Henrik Strahl (Newcastle) for communicating unpublished results prior to publication. We are also grateful to Anna Becker and Vivian Hennig (Kiel) for help in the initial stages of this project as part of their BSc theses.

## Author Contributions

M.B. and H.F. conceived the study, H.F. constructed the strains, performed the *in vivo* and *in vitro* experiments incl. microscopy. C.D. performed *in vitro* experiments. H.F., C.D. and M.B. analyzed the data, H.F. and M.B. wrote the manuscript.

## Declaration of Interests

The authors declare no competing interests.

## Appendix

### Plasmid and strain construction

All plasmids generated in this study were subjected to sequencing of relevant regions to verify plasmid integrity and to avoid occurrence of mutations. The oligonucleotides, plasmids and strains used in this study are listed in Supplementary files 1-3, respectively. *E. coli* NEB5-alpha was used to amplify and maintain plasmids.

### *E. coli* overexpression vector construction and transformation

For protein expression, the respective pET-28a plasmid variant was freshly transformed into BL21(DE3)/pLysS the day before expression using appropriate selection.

pET-28a-*minD* (NAP013). The *minD* gene was PCR amplified from *B. subtilis* 168 gDNA using primers NA018 and NA019. Subsequently, pET-28a (+) and the purified PCR product were digested using NdeI and XhoI and afterwards ligated and transformed.

pET-28a-*minD*-G12V (EHF045). This plasmid was generated using the Q5 Site-Directed Mutagenesis Kit (New England Biolabs) on plasmid NAP013 in combination with primers HF0213 and HF0214.

pET-28a-*minD*-K16A (EHF046). This plasmid was generated using the Q5 Site-Directed Mutagenesis Kit (New England Biolabs) on plasmid NAP013 in combination with primers HF0215 and HF0216.

pET-28a-*minD*-I260E (EHF047). This plasmid was generated using the Q5 Site-Directed Mutagenesis Kit (New England Biolabs) on plasmid NAP013 in combination with primers HF0237 and HF0238.

pET-28a-*minD*-D40A (EHF052). This plasmid was generated using the Q5 Site-Directed Mutagenesis Kit (New England Biolabs) on plasmid NAP013 in combination with primers HF0239 and HF0240.

pET-28a-*minC* (NAP012). The *minC* gene was PCR amplified from *B. subtilis* 168 gDNA using primers NA016 and NA017. Subsequently, pET-28a (+) and the purified PCR product were digested using NdeI and XhoI and afterwards ligated and transformed.

pET-28a-PDZ (CD001). The truncated *minJ* gene containing the PDZ-domain sequence was PCR amplified from *B. subtilis* 168 gDNA using primers HF0255 and HF0256. Subsequently, pET-28a (+) and the purified PCR product were digested using BamHI and XhoI and afterwards ligated and transformed.

### *B. subtilis* integration plasmid construction, transformation and selection of Halo-MinD constructs

Plasmid construction was verified via individual control digestion and DNA sequencing. Correct plasmids were transformed into *B. subtilis* strain 168 with the respectively indicated genetic background and selected for the introduced resistance. Resistant candidates were confirmed via PCR and microscopy.

pUC18-*minC-aad9-haloTag-minD-minDdown* (EHF054). This plasmid was constructed by assembling pUC18 (81) with three PCR products encoding *minC* (2), *aad9*-*haloTag* (3, encoding the spectinomycin adenyltransferase gene *aad9* and the *haloTag* gene) and *minD-minDdown* (4, encoding the *minD* gene and sequence downstream of *minD*) using the NEBuilder HiFi DNA Assembly Cloning Kit (New England Biolabs) according to protocol. The pUC18 plasmid was SmaI digested and purified, while the other three assembly pieces where PCR amplified. Specifically, the *minC* sequence with overhangs for pUC18 and *aad9* homology was amplified from gDNA template of 168 using primers HF0245 and HF0246. The *aad9-haloTag* sequence with overhangs for *minC* and *minD* homology was PCR amplified from plasmid template EHF042 using primers HF0247 and HF0248. The *minDdown* sequence with overhangs for *minD* and pUC18 homology was PCR amplified from from gDNA template of 168 using primers HF0249 and HF0250.

pUC18-*minC-aad9-haloTag-minD-G12V-minDdown* (EHF055). This plasmid was generated using the Q5 Site-Directed Mutagenesis Kit (New England Biolabs) on plasmid EHF054 in combination with primers HF0213 and HF0214.

pUC18-*minC-aad9-haloTag-minD-K16A-minDdown* (EHF056). This plasmid was generated using the Q5 Site-Directed Mutagenesis Kit (New England Biolabs) on plasmid EHF054 in combination with primers HF0215 and HF0216.

pUC18-*minC-aad9-haloTag-minD-D40A-minDdown* (EHF057). This plasmid was generated using the Q5 Site-Directed Mutagenesis Kit (New England Biolabs) on plasmid EHF054 in combination with primers HF0239 and HF0240.

pUC18-*minC-aad9-haloTag-minD-I260E-minDdown* (EHF058). This plasmid was generated using the Q5 Site-Directed Mutagenesis Kit (New England Biolabs) on plasmid EHF054 in combination with primers HF0237 and HF0238.

## Supplemental figures

**Figure 1–figure supplement 1:**
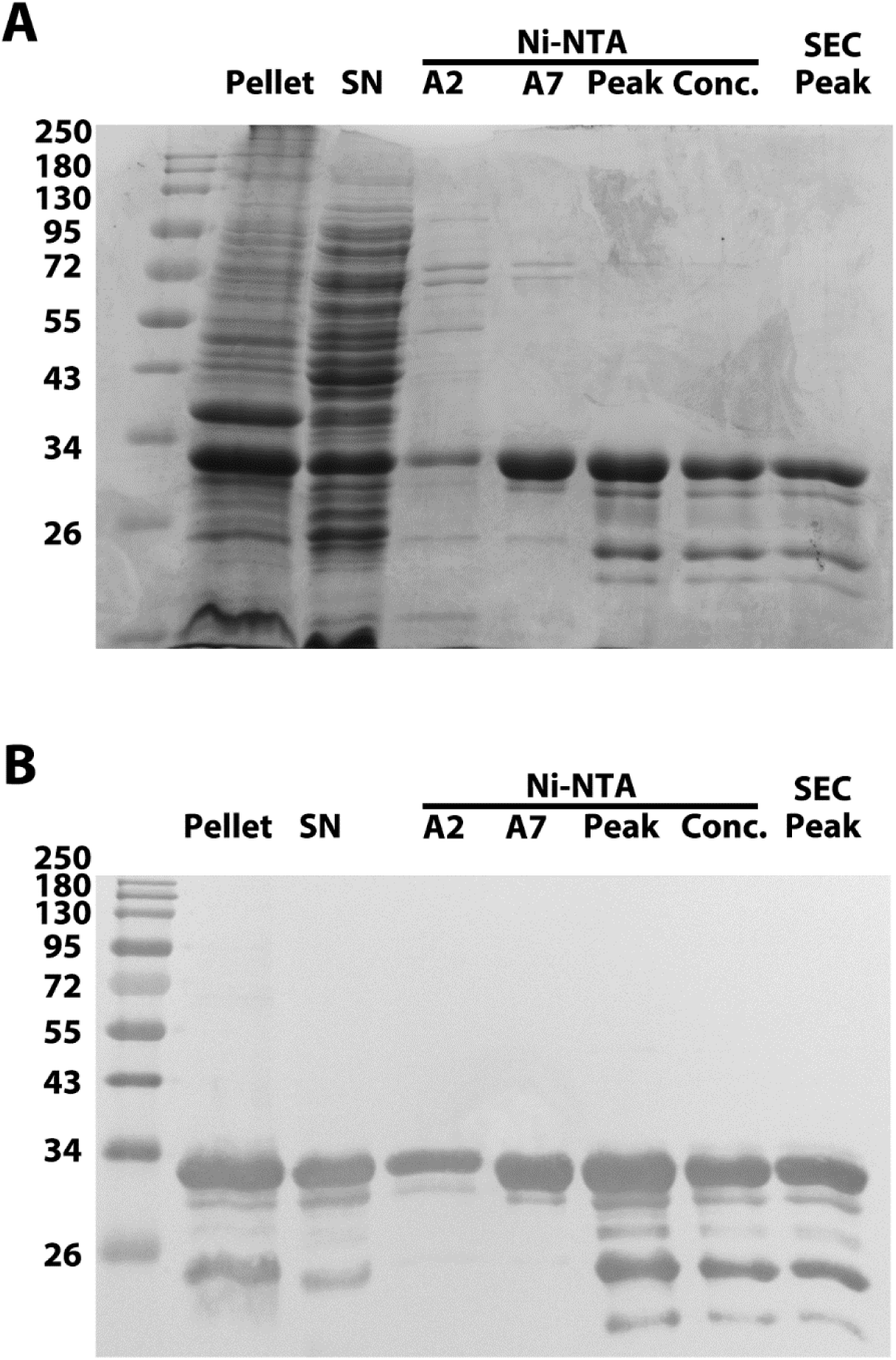
His-MinD purification analysis. (A) SDS-PAGE gel analysis of purification products of His-MinD (31.7 kDa), stained with Coomassie blue stain. From left to right: Ladder, pellet after lysate, supernatant, Ni-NTA peaks A2, A7, C4 (Main peak), concentrated sample, gel filtration peak. Ladder = Color Prestained Protein Standard, Broad Range (New England Biolabs). (B) Western blot of replicate gel from (A) using Penta-His antibody (QIAGEN).

**Figure 1–figure supplement 2:**
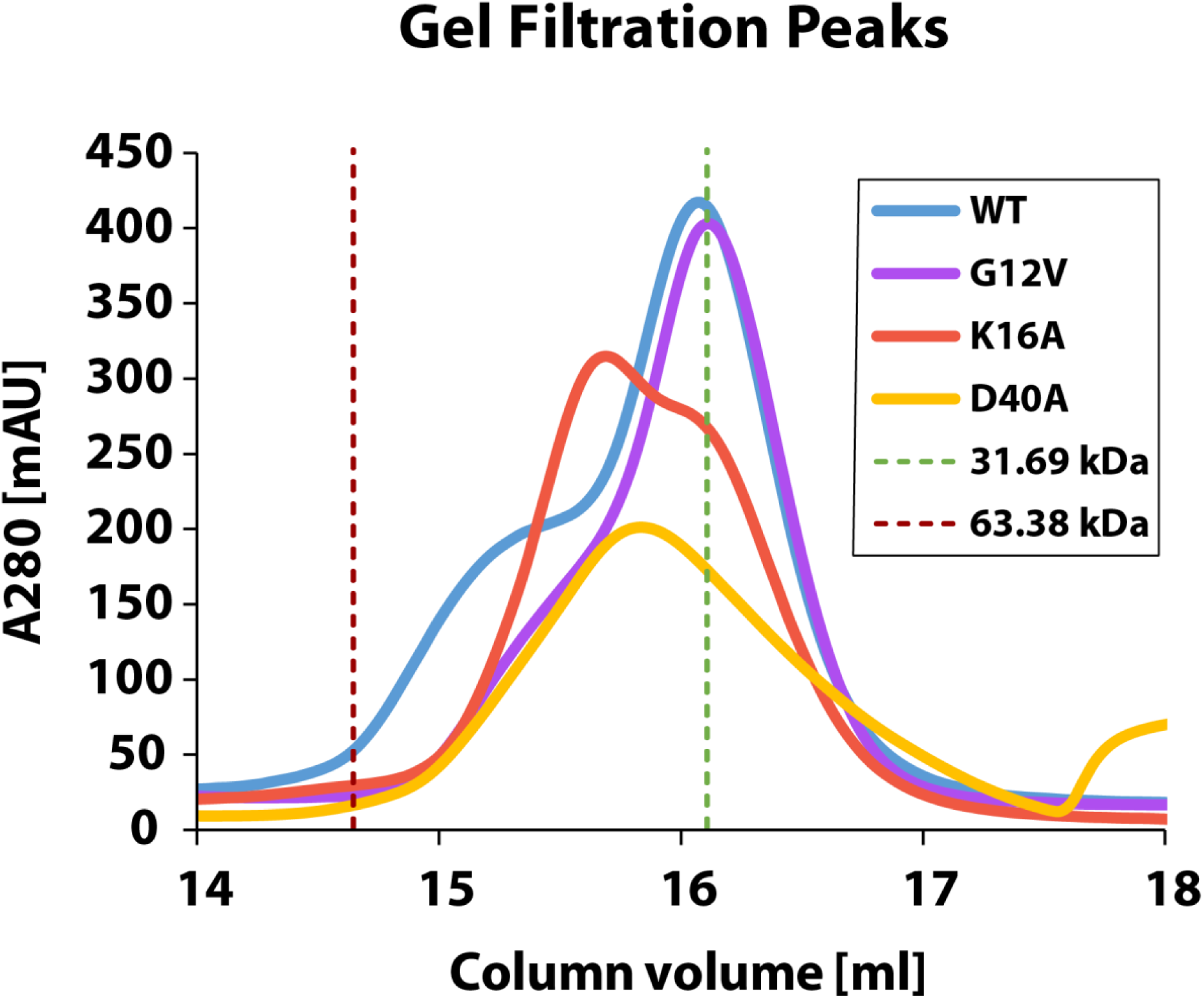
Gel filtration chromatograms of His-MinD and variants. Gel filtration analysis of purified His-MinD and indicated mutants. Curves were aligned according to the respective void volume after sample injection. Green and red dotted lines mark the calibrated elution volumes corresponding to proteins of 31.69 kDa (His-MinD monomer) and 63.38 kDa (His-MinD dimer), as determined from molecular-weight standards.

**Figure 2–figure supplement 1:**
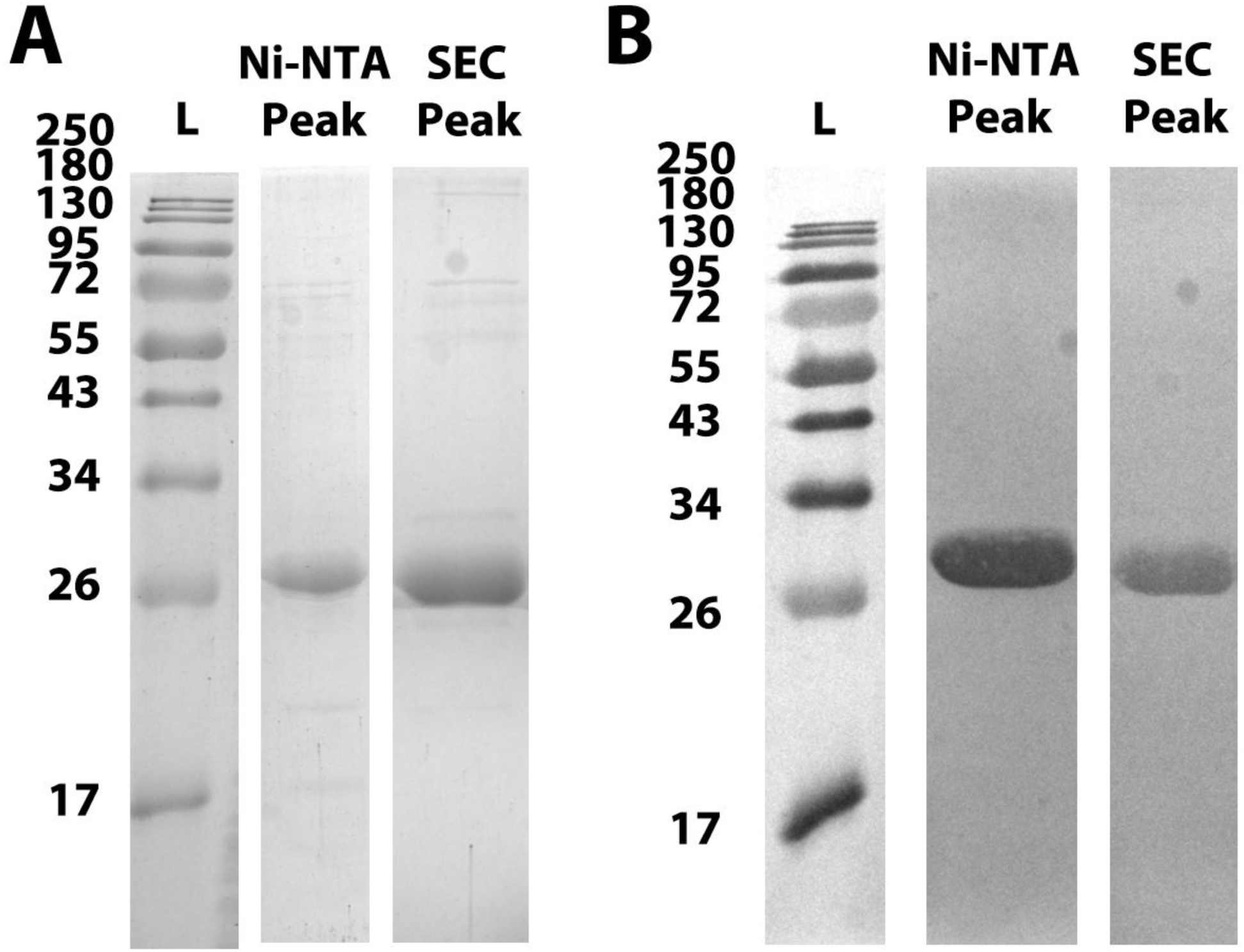
His-MinC purification analysis. (A) SDS-PAGE gel analysis of purification products of His-MinC (27.3 kDa), stained with Coomassie blue stain. From left to right: Ladder, Ni-NTA peak, SEC peak. Ladder = Color Prestained Protein Standard, Broad Range (New England Biolabs). (B) Western blot of replicate gels from (A) using Penta-His antibody (QIAGEN).

**Figure 2–figure supplement 2:**
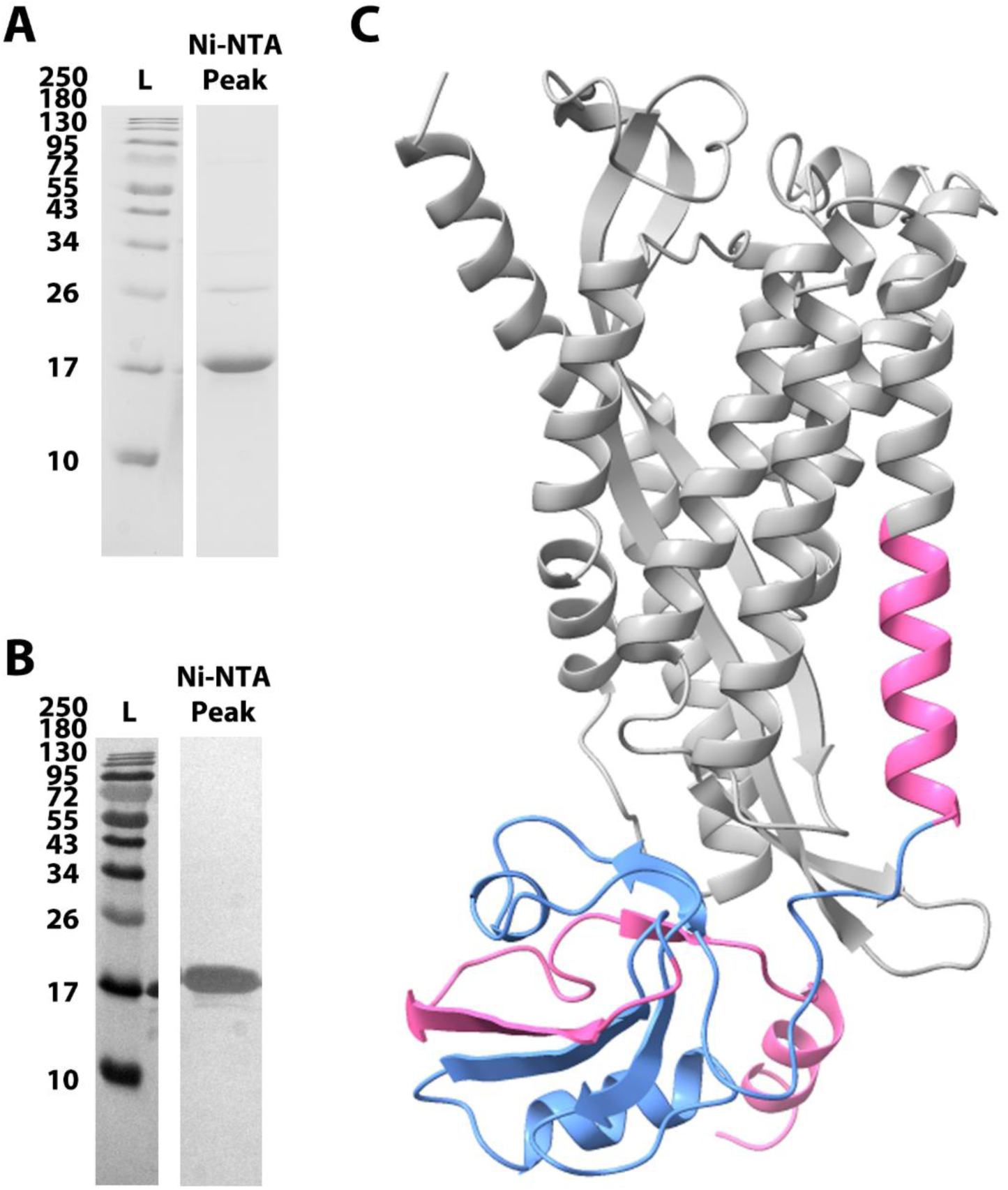
His-PDZ purification analysis and rendering of MinJ AlphaFold model. (A) SDS-PAGE gel analysis of purification products of His-PDZ (17.6 kDa), stained with Coomassie blue stain. From left to right: Ladder (L), Ni-NTA main peak. Ladder = Color Prestained Protein Standard, Broad Range (New England Biolabs). (B) Western blot of replicate gel from (A) using Penta-His antibody (QIAGEN). (C) Rendering of AlphaFold 3 model of *B. subtilis* MinJ (grey) (83). The truncation expressed with pET-28a-PDZ, which includes the PDZ domain (blue), is highlighted in pink.

**Figure 4–figure supplement 1:**
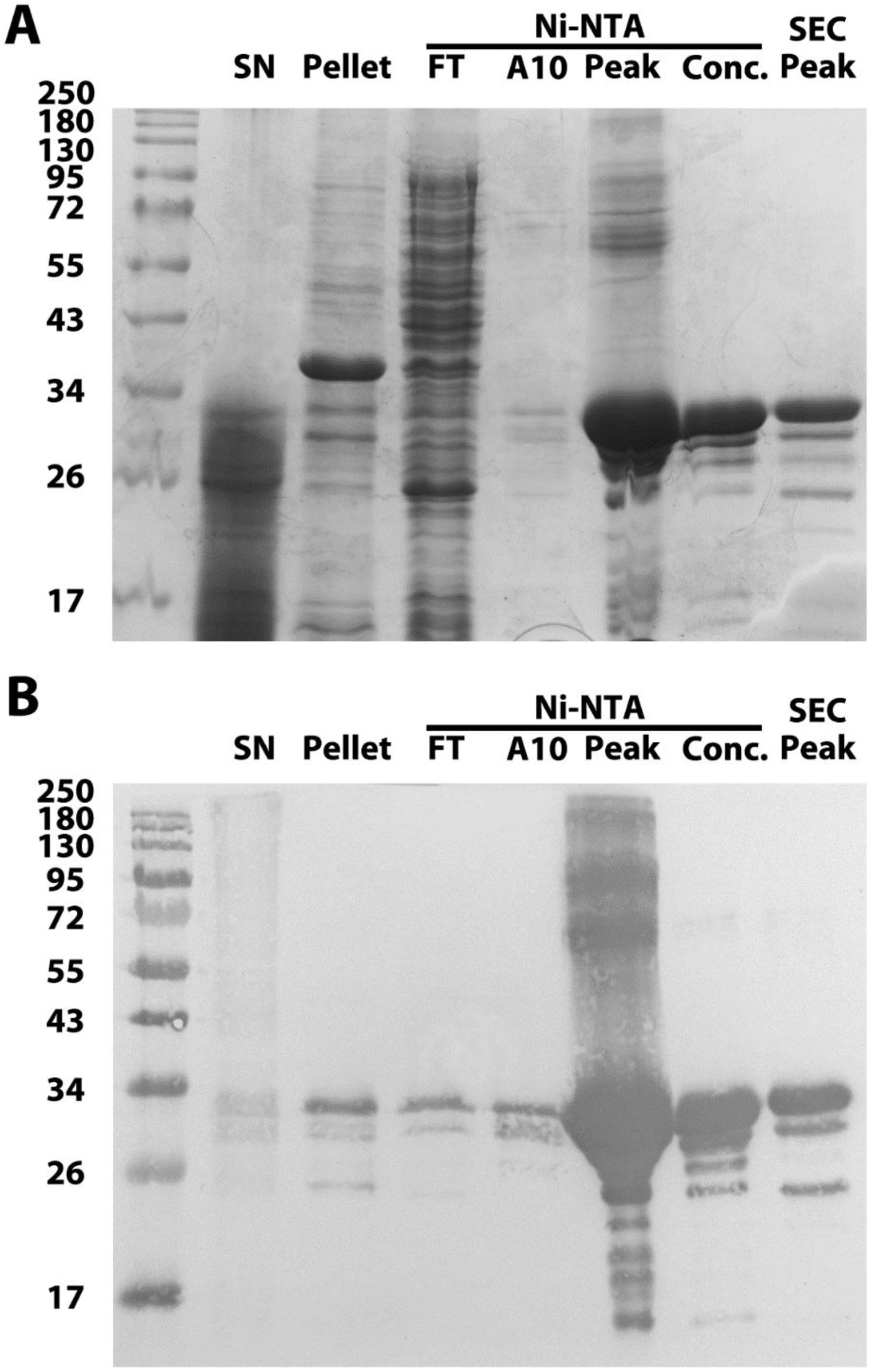
His-MinD-I260E purification analysis. (A) SDS-PAGE gel analysis of purification products of His-MinD-I260E (31.7 kDa), stained with Coomassie blue stain. From left to right: Ladder, supernatant after lysis, pellet, Ni-NTA flowthrough, Ni-NTA peaks A10 and C4 (Main peak), concentrated sample, gel filtration peak. Ladder = Color Prestained Protein Standard, Broad Range (New England Biolabs). (B) Western blot of replicate gel from (A) using Penta-His antibody (QIAGEN).

**Figure 4–figure supplement 2:**
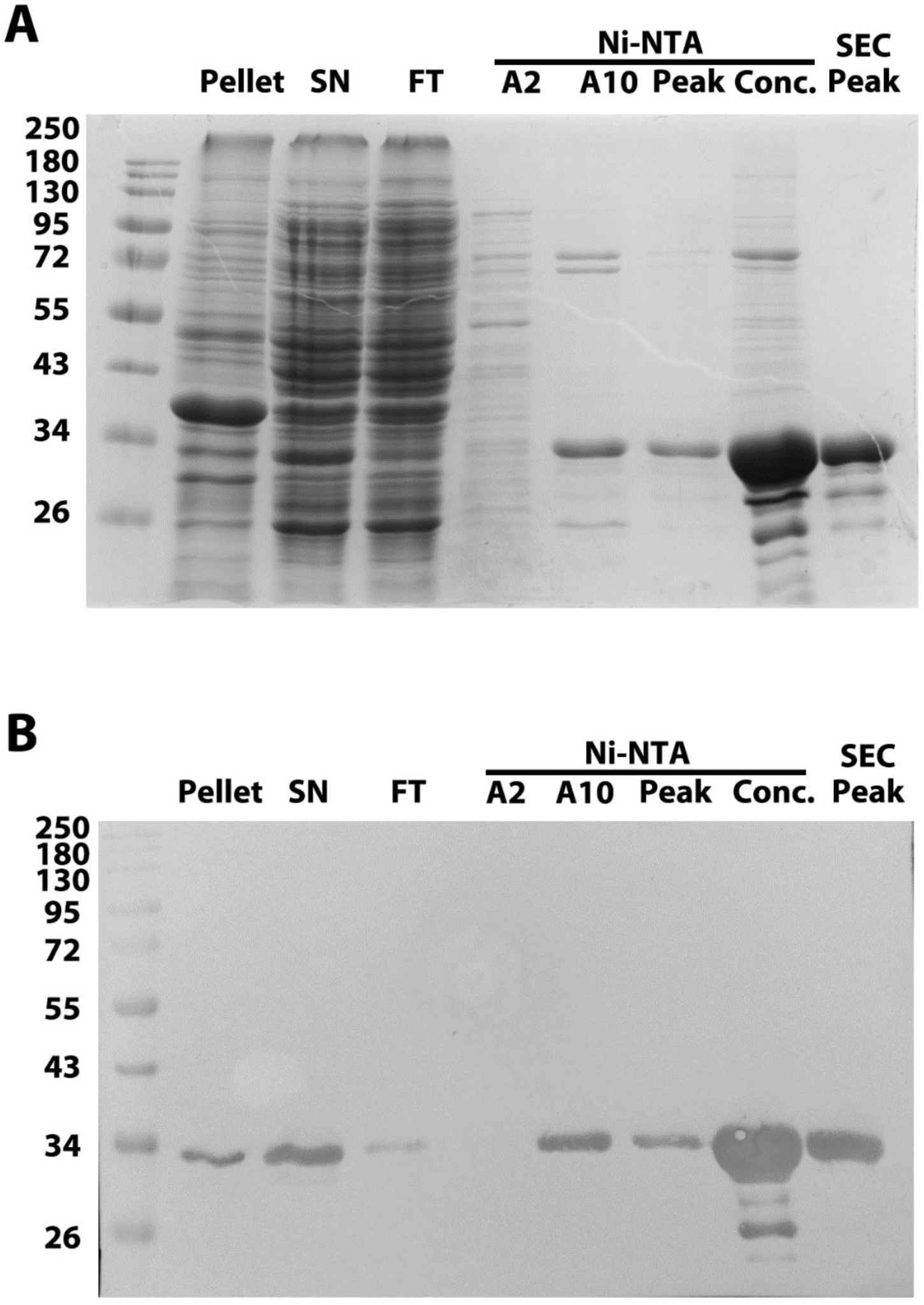
His-MinD-G12V purification analysis. (A) SDS-PAGE gel analysis of purification products of His-MinD-G12V (31.7 kDa), stained with Coomassie blue stain. From left to right: Ladder, pellet after lysate, supernatant, Ni-NTA flowthrough, Ni-NTA peaks A2, A10, C4 (Main peak), concentrated sample, gel filtration peak. Ladder = Color Prestained Protein Standard, Broad Range (New England Biolabs). (B) Western blot of replicate gel from (A) using Penta-His antibody (QIAGEN).

**Figure 4–figure supplement 3:**
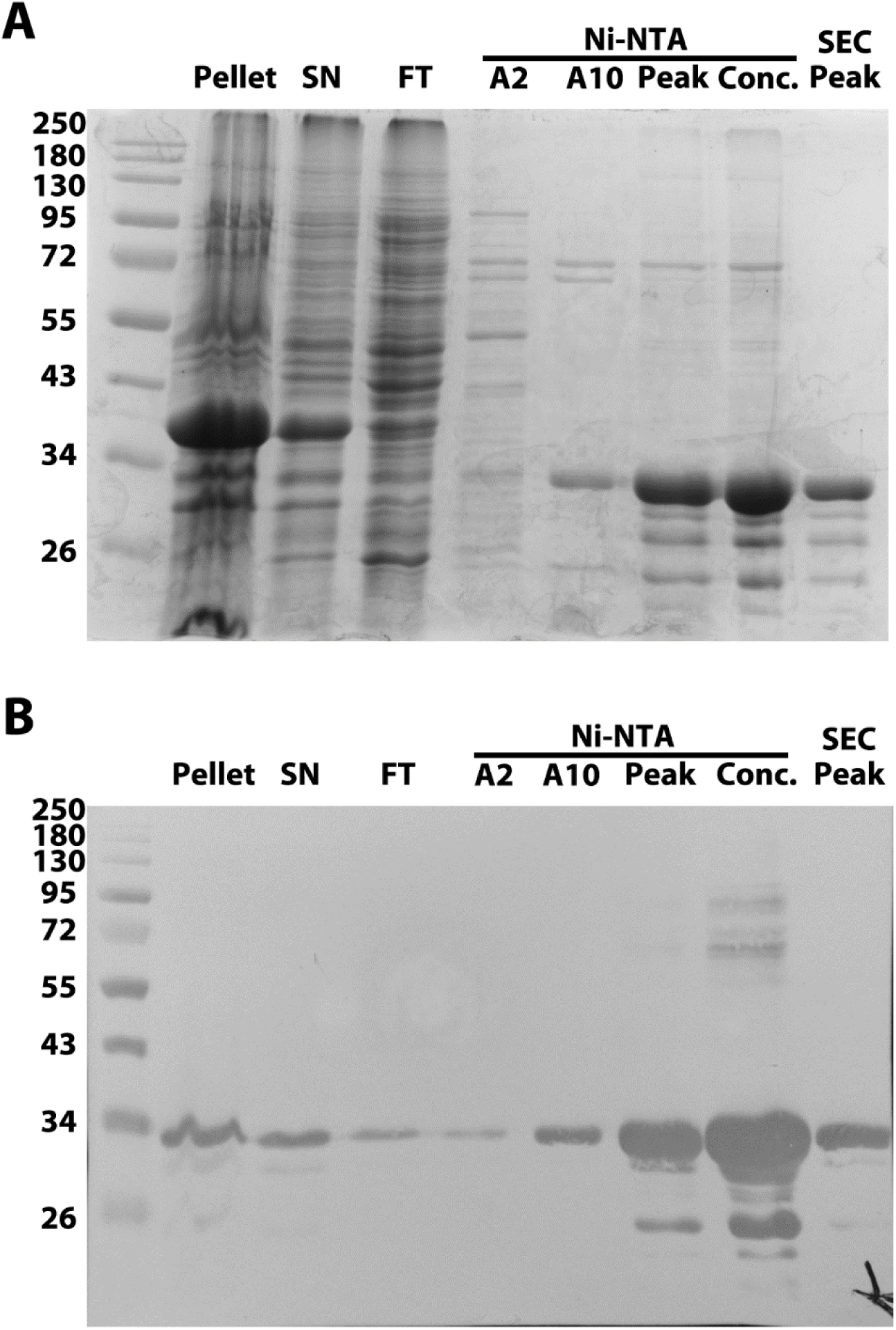
His-MinD-K16A purification analysis. A) SDS-PAGE gel analysis of purification products of His-MinD-K16A (31.7 kDa), stained with Coomassie blue stain. From left to right: Ladder, pellet after lysate, supernatant, Ni-NTA flowthrough, Ni-NTA peaks A2, A10, C4 (Main peak), concentrated sample, gel filtration peak. Ladder = Color Prestained Protein Standard, Broad Range (New England Biolabs). (B) Western blot of replicate gel from (A) using Penta-His antibody (QIAGEN).

**Figure 4–figure supplement 4:**
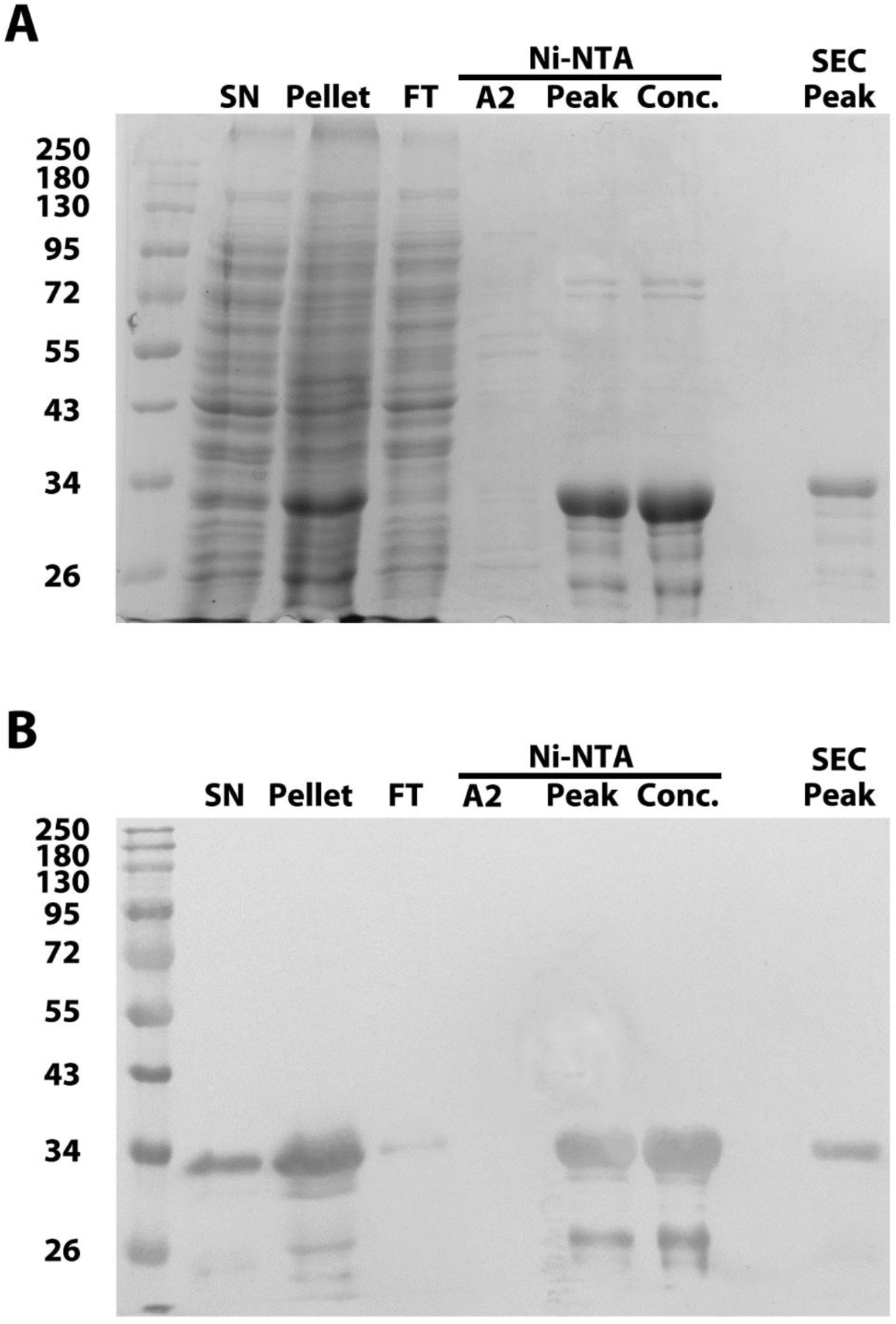
His-MinD-D40A purification analysis. (A) SDS-PAGE gel analysis of purification products of His-MinD-D40A (31.7 kDa), stained with Coomassie blue stain. From left to right: Ladder, supernatant after lysis, pellet, Ni-NTA flowthrough, Ni-NTA peaks A2 and C4 (Main peak), concentrated sample, gel filtration peak. Ladder = Color Prestained Protein Standard, Broad Range (New England Biolabs). (B) Western blot of replicate gel from (A) using Penta-His antibody (QIAGEN).

**Figure 5–figure supplement 1:**
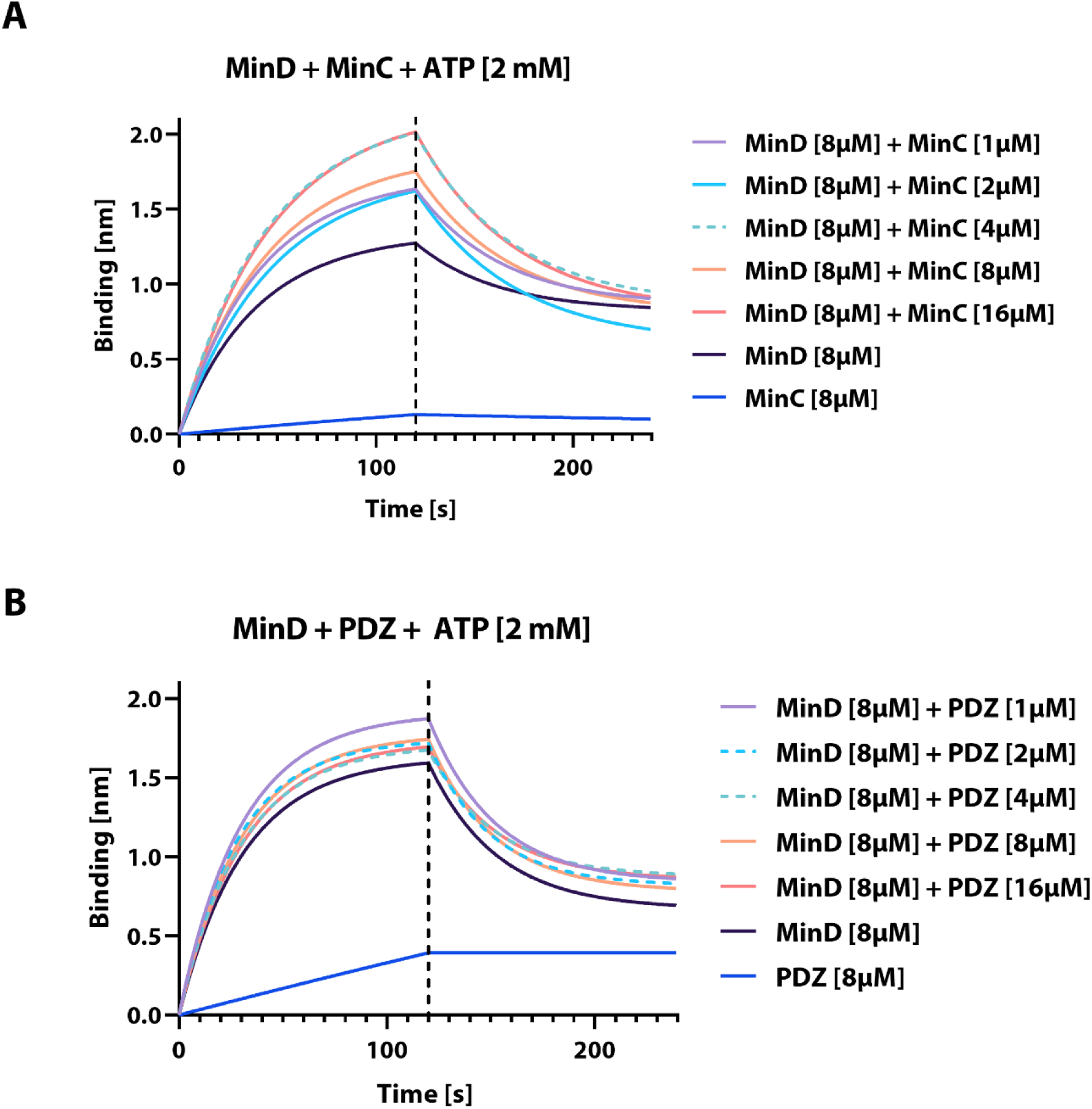
BLI analysis of interactions between His-MinD and His-MinC or His-PDZ. (A) MinD binding (see Figure 5 A) plotted against time through association and dissociation phase (steps 4-5) at different indicated His-MinC concentrations, including His-MinD and His-MinC only controls, all in presence of freshly added ATP [2 mM]. (B) Same as (A) using the indicated respective concentrations of His-PDZ.

**Figure 5–figure supplement 2:**
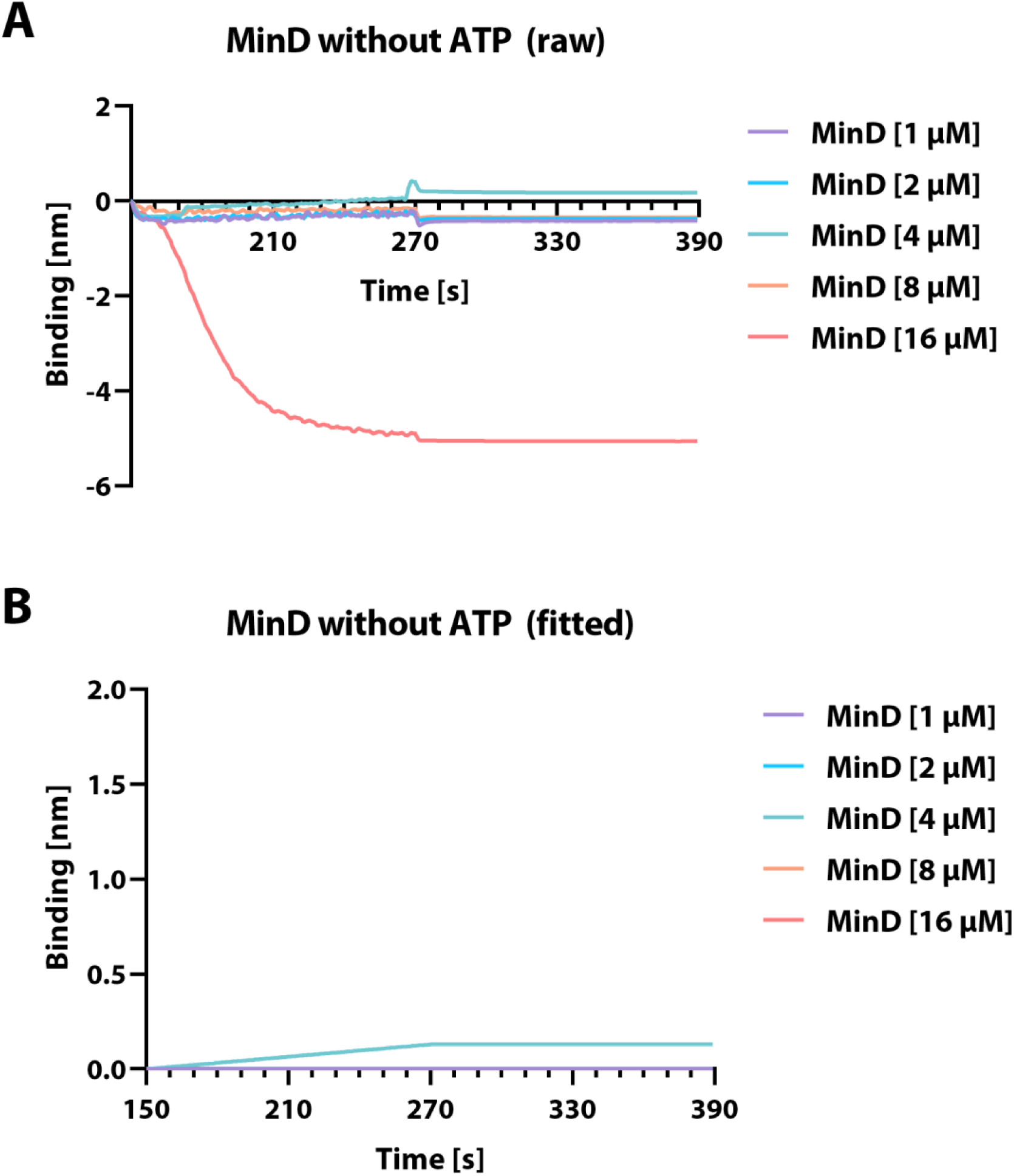
BLI analysis of His-MinD without addition of ATP. His-MinD binding to liposome associated BLI sensor (see Figure 5), plotted against time through association and dissociation phase at different protein concentrations without the addition of Mg^2+^-ATP.

## Supplemental tables

**Supplementary table 1:**
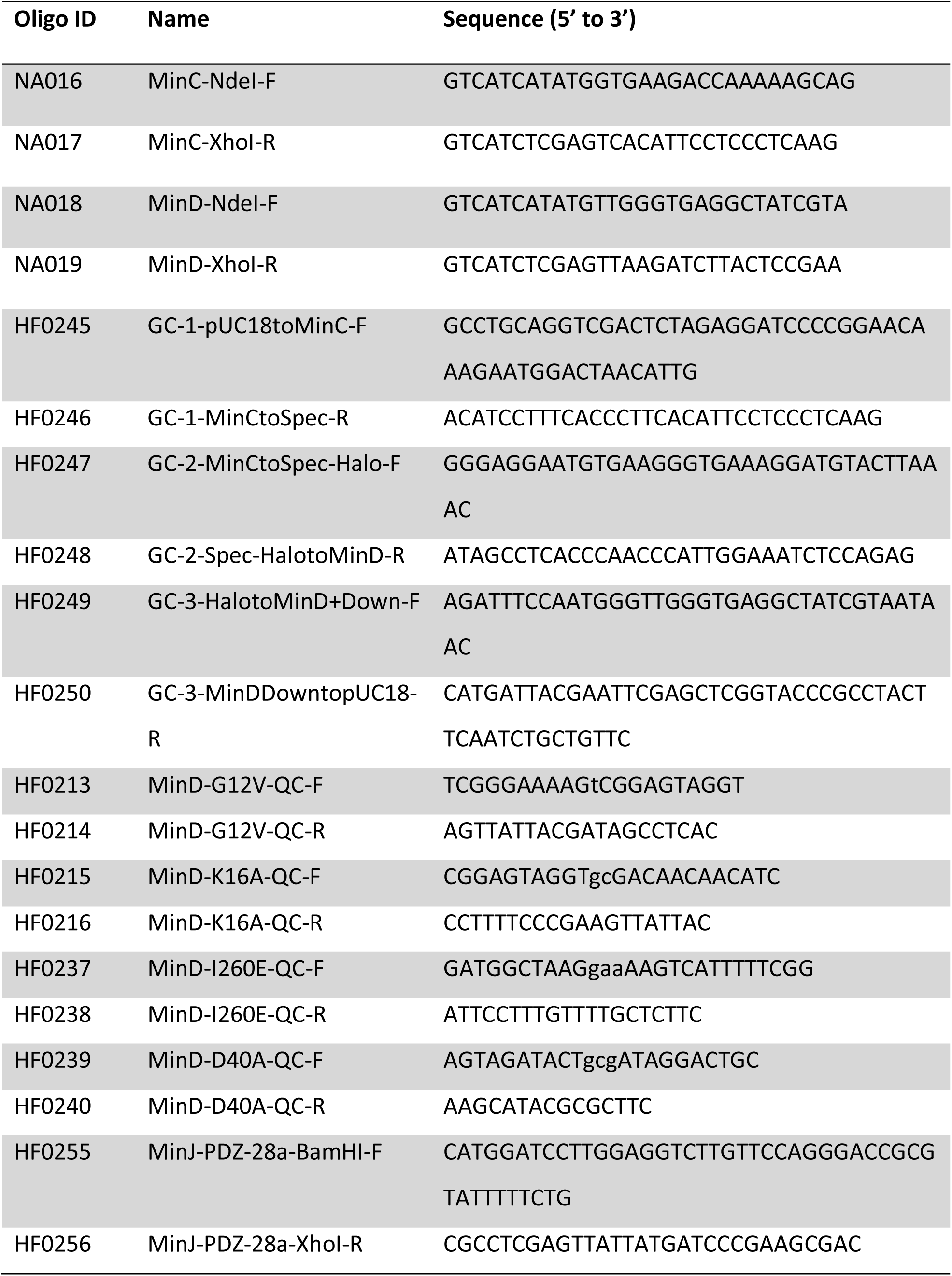
Oligonucleotides used in this study.

**Supplementary table 2:**
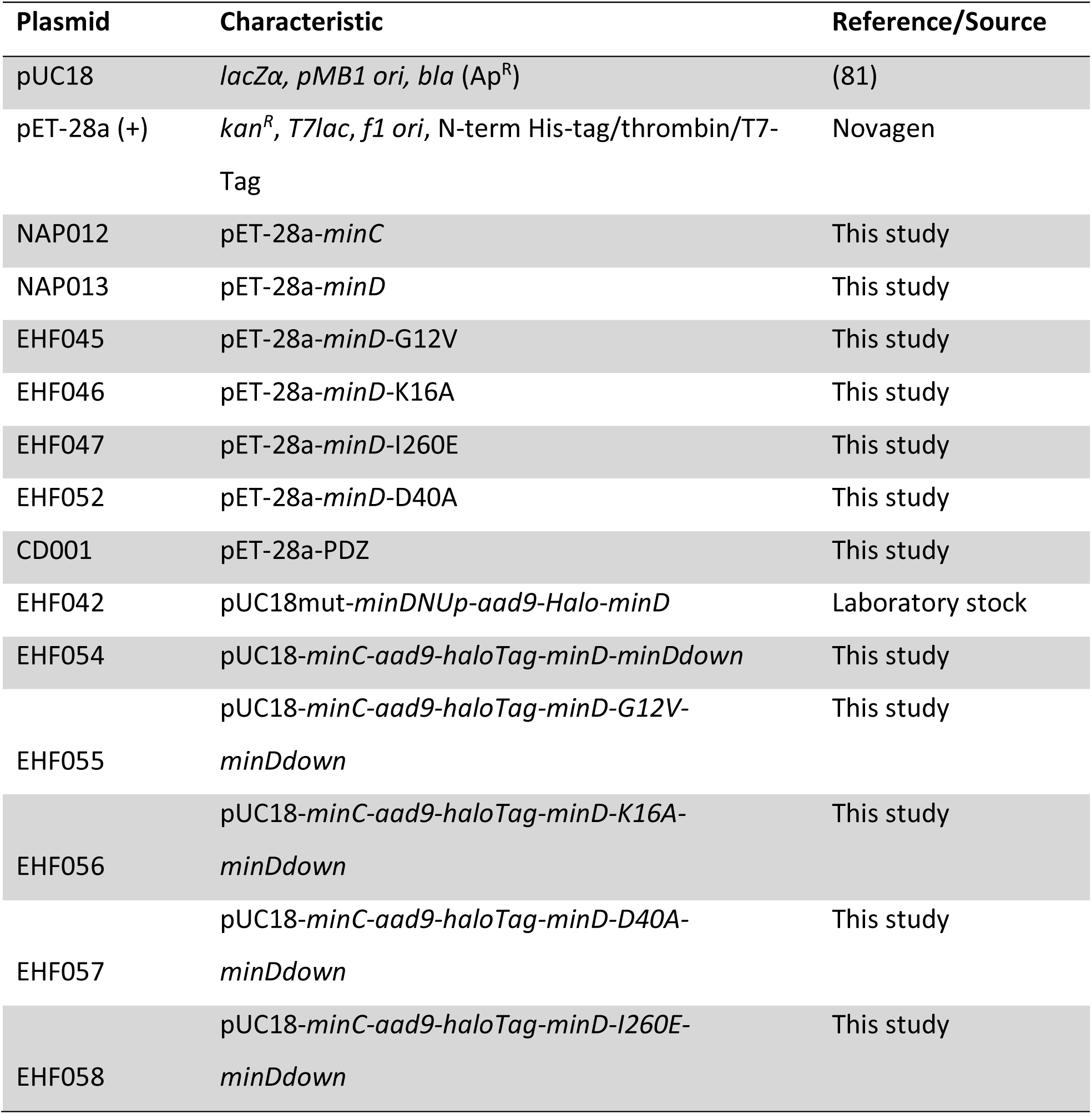
Plasmids used in this study.

**Supplementary table 3:** Strains used in this study.

| Strain | Relevant features/genotype | Reference/Source | Transformation |
| --- | --- | --- | --- |
| ***B. subtilis*** |  |  |  |
| 168 | *trpC2* | Laboratory collection |  |
| BKK28000 | *minC::lox71-kanR-lox66* | Addgene (*B. subtilis*<br>BKK Library, Kan, (97)) |  |
| RD021 | *minJ::tet* | (39) |  |
| BKK27990 | *minD::lox71-kanR-lox66* | Addgene (*B. subtilis*<br>BKK Library, Kan, (97)) |  |
| BHF077 | *minD::lox71-kanR-lox66* | This study | BKK27990 gDNA<br>-> 168 |
| BHF078 | *minD::aad9-Halo-minD* | This study | EHF054 -><br>BHF077 |
| BHF079 | *minD::aad9-Halo-minD-G12V* | This study | EHF055 -><br>BHF077 |
| BHF080 | *minD::aad9-Halo-minD-K16A* | This study | EHF056 -><br>BHF077 |
| BHF081 | *minD::aad9-Halo-minD-D40A* | This study | EHF057 -><br>BHF077 |
| BHF082 | *minD::aad9-Halo-minD-I260E* | This study | EHF058 -><br>BHF077 |
| BHF083 | *minD::aad9-Halo-minD,*<br>*minC::lox71-kanR-lox66* | This study | BKK28000 gDNA<br>-> BHF078 |
| BHF084 | *minD::aad9-Halo-minD, minJ::tet* | This study | RD021 gDNA -><br>BHF078 |
| ***E. coli*** |  |  |  |
| NEB 5-alpha | *fhuA2Δ(argF-lacZ)U169 phoA*<br>*glnV44 Φ80Δ(lacZ)M15 gyrA96*<br>*recA1 relA1 endA1 thi-1 hsdR17* | New England Biolabs |  |
| BL21(DE3)pLysS | F<sup>-</sup>, *ompT*, *hsdS<sub>B</sub>* (r<sub>B</sub><sup>-</sup>, m<sub>B</sub><sup>-</sup>), *dcm*,<br>*gal*, λ(DE3), pLysS, Cm<sup>r</sup> | Promega |  |

**Supplementary table 4:**
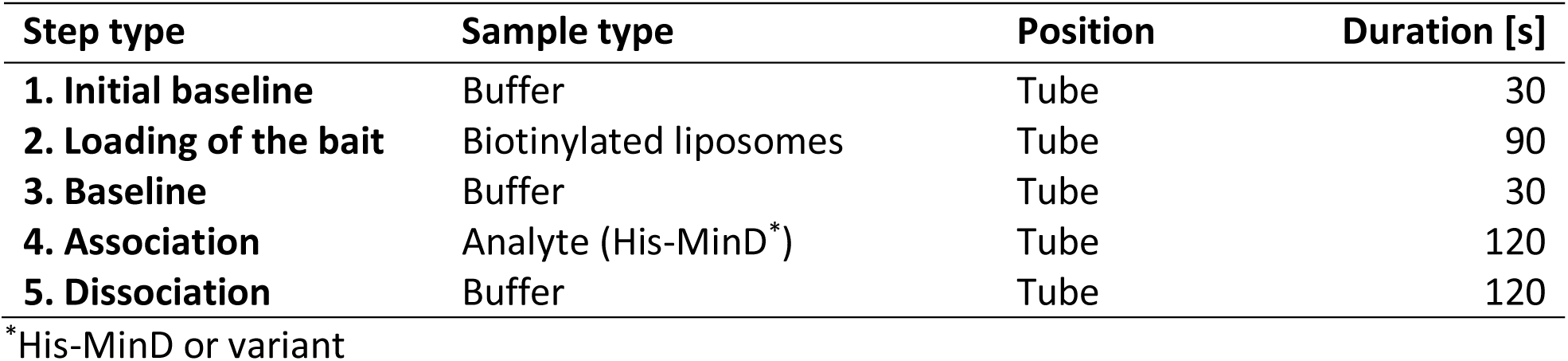
BLI Advanced Kinetics Protocol.

| Step type | Sample type | Position | Duration [s] |
| --- | --- | --- | --- |
| **1. Initial baseline** | Buffer | Tube | 30 |
| **2. Loading of the bait** | Biotinylated liposomes | Tube | 90 |
| **3. Baseline** | Buffer | Tube | 30 |
| **4. Association** | Analyte (His-MinD*) | Tube | 120 |
| **5. Dissociation** | Buffer | Tube | 120 |

\*His-MinD or variant

## Notes

### Competing Interest Statement

The authors have declared no competing interest.

### Summary of Updates

This revised version contains additional sections in the discussion and has a rearranged supllement.

